# Evolutionary basis of intermale sexual behavior by multiple pheromone switches in *Drosophila*

**DOI:** 10.1101/2025.10.14.682417

**Authors:** Youcef Ouadah, Thomas H. Naragon, Hayley Smihula, Emily L. Behrman, Mohammed A. Khallaf, Yun Ding, David L. Stern, Joseph Parker, David J. Anderson

**Author notes:** Department of Biological Sciences, Dartmouth College; Hanover, NH, USA. Molecular Physiology of Somatic Sensation Laboratory, Max Delbrück Center for Molecular Medicine; Berlin, Germany and Department of Zoology and Entomology, Faculty of Science, Assiut University; Assiut, Egypt (Mohammed A. Khallaf). Stowers Institute for Medical Research and Howard Hughes Medical Institute; Kansas City, MO, USA.

## Abstract

We have identified a *Drosophila* species which exhibits spontaneous and robust intermale sexual behavior. *D. santomea* males distinguish conspecific sexes but court both vigorously and seldom attack. Elevated intermale courtship stems from at least three evolutionarily derived pheromonal changes. In males, the sexually monomorphic cuticular pheromone 7-tricosene promotes rather than inhibits courtship and the courtship-inhibiting olfactory pheromone cVA is reduced 84-92% compared to close relatives. The third switch is in *D. santomea* females, where cVA suppresses rather than promotes sexual receptivity. Female cVA aversion and male cVA reduction may have co-evolved to maintain efficient intraspecific mating but prevent hybridization with the sympatric sibling species D*. yakuba*. High intermale courtship and low cVA also co-occur and appear selectively derived in a distant monomorphic species *D. persimilis*, implying pheromonal and social behavioral convergence. Changes in pheromone valence and levels may therefore explain the recent evolutionary emergence of intermale sexual behavior in *Drosophila*.

## Introduction

Social behaviors, including aggregation, aggression, courtship, mating, parenting, and cooperation, are remarkably diverse in the animal kingdom (*1*), and can vary between closely related sibling species (*2*) or even entire orders (*3*). The ultimate causes of social behaviors (in terms of natural and sexual selection) have received considerable theoretical attention (*4*, *5*) and proximate causes (including genetic loci, hormones, neural circuits, pheromones, and sensory releasers) identified (*6–15*). The ∼2,000 species of the genus *Drosophila* (*16*) have historically excelled in this regard as models for intersexual social behaviors (*17–22*), providing notably an unparalleled genetic, neural, and pheromonal understanding of the iconic courtship dance of *D. melanogaster* males (*23–25*). Comparisons of male courtship in other members of the genus have revealed flexible substrates, in genomes (*26–29*) and nervous systems (*30–33*), of its rapid evolution and revealed general principles of reproductive function and isolation (*34*, *35*). However, comparative investigations of intrasexual social behaviors between *Drosophila* males or females have relatively lagged.

Interactions between same-sex conspecifics can vary greatly across species, strains, individuals, and time (*36*, *37*), and can be as opposed as altruistic cooperation (*38*) or lethal aggression (*39*). Such flexibility implies that intrasexual social behaviors can evolve quickly under varying selection pressures or neutral drift, but the underlying changes in pheromone systems, neural circuits, and genomes are often difficult to pinpoint. We sought to identify phylogenetic patterns of variation in intermale social behavior in *Drosophila* and, by leveraging experimental strengths of the genus, reveal proximate and ultimate evolutionary causes. We identified a species, *D. santomea*, in which males show vigorous sexual behavior toward other males that includes multiple species-typical motor and acoustic features of the female-directed courtship dance, and is accompanied only rarely by overt aggression. Three distinct changes in pheromone levels or behavioral valence compared to *D. melanogaster* have occurred during *D. santomea* evolution, both before and after divergence from the sibling species *D. yakuba*. Together these changes have highly and selectively amplified intermale courtship without eliminating sex discrimination or the latent potential for intense attack, which can be revealed by optogenetic activation of conserved aggression-promoting neural circuits. The results, together with information from natural history and phylogenetic inference, provide insight into the pheromonal mechanism and evolutionary logic of recent social evolution in *Drosophila* that may apply broadly to other taxa.

## Results

### Evolutionary variation in male-male social behavior within the genus *Drosophila*

In the presence of food or a female, pairs of single-housed *D. melanogaster* males will fight (*40–48*), employing a stereotyped set of contact- and non-contact-mediated motor actions including lunging, tussling, and wing threat (*49*, *50*). We assessed aggression (and other intermale social behaviors) comparatively in 15 *Drosophila* species using food as the stimulating resource (*51*), screening for qualitative and quantitative variation in male social engagement, frequency, and motor pattern. We sampled species densely within the *melanogaster* subgroup (7 tested of 9 assigned) (*52*) to monitor potential behavior variation at close phylogenetic distances to *D. melanogaster* with high resolution. At larger evolutionary distances we sampled sparsely (typically one species per subgenus group or subgroup) to broadly survey genus social evolution, apart from including both members of the *D. pseudoobscura* / *D. persimilis* sibling species pair from the *obscura* group (*53*). We standardized assay conditions but occasionally varied the food source over which flies competed to account for host specializations (*54*). Fly behavior was measured in dyadic male-male interactions by combining saturated manual scoring of 11 social actions on a subset of representative recordings (Fig. 1A,B, fig. S1A) with automated behavior classifications for the remainder (fig. S2A,B) (*55–57*). Separate behavior classifiers detected either lunge, a major consummatory attack action in males (*43*, *49*, *50*), or unilateral wing extension (UWE), a cross-genus indicator of male sexual behavior (*20*), as proxies for aggression and courtship, respectively.

**Figure 1.**
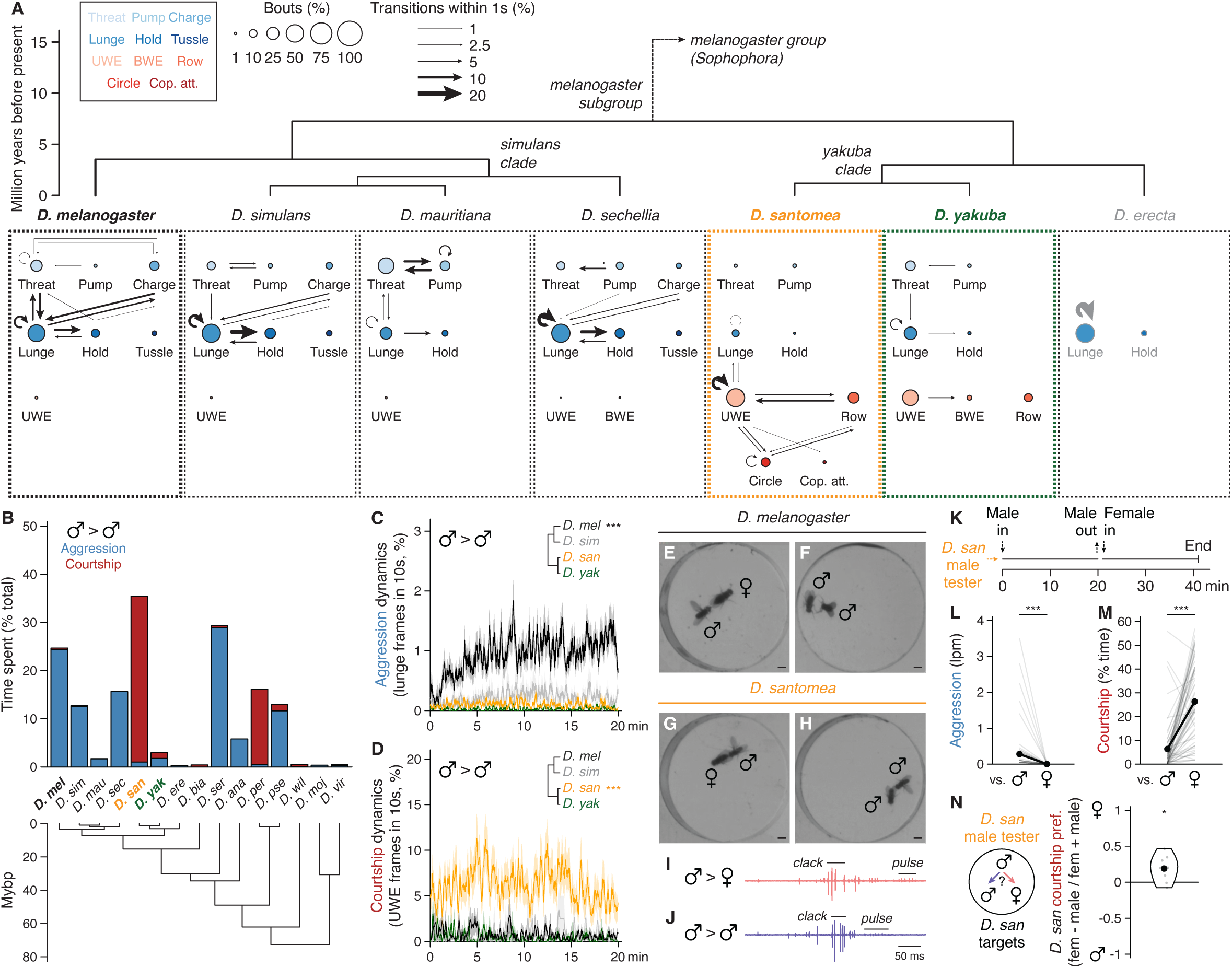
Evolutionary variation in male-male social behavior within the genus *Drosophila*. **(A)** Ethograms from twenty-minute male-male interactions over food depicting aggressive and courtship actions scored in three representative conspecific pairs per species across the melanogaster subgroup. Nodes are bout counts of a given action type, with sizes representing frequency normalized to summed bouts scored across all actions (expert annotation). Edges indicate action transitions, with weights representing the fraction of bouts of a given action (arrow origin) for which a second action (arrow destination) occurred within one second. Eleven total actions in each ethogram are arranged in four rows: first, three aggressive threat actions (threat, pump, charge) (*106*); second, three aggressive contact-mediated actions (lunge, hold, tussle) (*49*); third, three courtship actions utilizing the wings and typically directed toward females (UWE, BWE, row) (*20*); fourth, two courtship actions involving locomotory maneuvers (circle, copulation attempt). Nodes indicating aggressive actions are filled with blue shades and courtship actions with reds. Species arrangement and divergence times according to compilation of available phylogenies (*75*, *154*, *155*). Note emergence of male-directed courtship and reduction of aggression in *D. santomea* (orange) and, to a lesser extent, its sympatric sister species *D. yakuba* (green). *D. erecta* ethogram gray coloration indicates that very few social actions of any kind were observed. Abbreviations: UWE, unilateral wing extension; BWE, bilateral wing extension; Cop. att., copulation attempt. See table S2 for wildtype strains used in figures throughout. **(B)** Stacked barplots showing cumulative time spent by paired males in any aggressive (blue) or courtship action (red) as a fraction of the twenty-minute total interaction. Additional Sophophora subgenus species for which male-male interactions were scored for the same action set are included. Note unusually high time dedication to male-directed courtship by *D. santomea. D. serrata* aggression scoring is inflated due to unusually long holds. **(C,D)** Expressivity and temporal dynamics of male-male aggression (C) and courtship (D) in *D. melanogaster* (black), *D. simulans* (gray), *D. santomea* (orange), and *D. yakuba* (green). Bold traces indicate the mean fraction of frames annotated as lunge (for aggression, C) or UWE (for courtship, D) by automated classifiers in a 10-second sliding window. Envelopes represent standard error of the mean (s.e.m.). Note high aggression in *D. melanogaster* male-male pairs and high courtship in *D. santomea* throughout twenty-minute interactions (significance by Kolmogorov-Smirnov tests). See table S4 for statistics related to figures throughout. **(E-H)** Video stills of *D. melanogaster* (E,F) and *D. santomea* males (G,H) interacting with conspecific female (E,G) or male targets (F,H). *D. melanogaster* court females (E, male shown in UWE) and attack males (F, shown mid-lunge). *D. santomea* court both sexes using a similar set of motor patterns (G,H, males shown in UWE for both) and rarely attack. Scale bars, 1 mm. **(I,J)** Representative *D. santomea* courtship song traces showing “pulse” and “clack” emitted toward both females (I) and males (J). **(K)** Scheme for assessing social behavioral differences by *D. santomea* male tester flies paired consecutively with a male then female conspecific target for twenty minutes each. **(L.M)** Aggression (L, measured as lunges per minute) and courtship (M, fraction of total interaction time performing UWE) exhibited by *D. santomea* male testers towards male then female targets in sequential pairings. Data points derived from the same tester are paired and distribution means shown as paired black dots. Note aggression towards males is never also observed to females and significantly increased courtship rates to females (paired Mann-Whitney *U* tests). **(N)** Courtship preference index exhibited by *D. santomea* male testers towards conspecific female and male targets presented simultaneously in ten-minute trio assays. Index calculated as the difference in time spent courting each sex normalized by the sum. Significance compared to a null median of zero (indicating no preference) determined by one-sample Mann-Whitney *U* test. Note slight but significant preference for courting females.

Aggression among *D. melanogaster* males was the most prominent and intense of any *melanogaster* subgroup species tested (Fig. 1A-C, fig. S2C,E). Species in the neighboring *simulans* clade (*D. simulans*, *D. mauritiana*, and *D. sechellia*) also fought, though less frequently, using a similar set of actions (Fig. 1A-C, fig. S2C,E). Varying foods usually had little effect on the behavior of specialists, with the notable exception of *D. sechellia* which fought significantly more over *Morinda* than the standard apple juice (fig. S3A), consistent with its specialization (*54*). Male-male interactions in *D. melanogaster* and *simulans* clade species all included very little recognizable courtship, and the few bouts we observed never progressed to attempted copulations (Fig. 1A,B). Thus, intermale aggressive actions in the subgroup branch that includes *D. melanogaster* are almost entirely distinct from the courtship that males perform toward females (Fig. 1E,F, fig. S4).

The other branch of the *melanogaster* subgroup, which includes *D. erecta*, *D. yakuba*, and *D. santomea*, exhibited dramatically different male-male dynamics. *D. erecta* males hardly interacted or locomoted, even on a *Pandanus* substrate (Fig. 1A,B, fig. S1B, fig. S3) (*58*). *D. yakuba* and *D. santomea,* by contrast, exhibited low levels of aggression and multiple distinct, clearly recognizable courtship actions (Fig. 1A,B). Male-male courtship in both species was visually similar as toward females (Fig. 1G,H, fig. S4), but the two sibling species differed markedly from each other in penetrance and expressivity. Courtship and attack between *D. yakuba* males were both infrequent, whereas *D. santomea* males courted other males considerably more than they fought them and far more than did *D. yakuba* or any other subgroup species (Fig. 1B,D, fig. S2D,F). High levels of male-directed courtship and low aggression were conserved in three additional *D. santomea* strains (fig. S2C-F), were not limited to the initial moments of an encounter when sex identification might be incomplete (Fig. 1D) (*51*, *59*), and sometimes culminated in copulation attempts (Fig. 1A). Audio recordings (*60*) confirmed that *D. santomea* males sang to males as they did to females, emitting both “pulse” and “clack” song (*61–63*) with similar (though not identical) acoustic features (Fig. 1I,J, fig. S5).

Thus, under our conditions we find a striking difference in intermale social behavior between *D. melanogaster* and *D. santomea*. *D. melanogaster* and *simulans* clade relatives attack males with varying frequencies using a conserved set of actions, whereas *D. santomea* courts both males and females with nearly indistinguishable motor and acoustic features. While widely described in other taxa (*64*) and various *D. melanogaster* mutants (*24*), the frequent courtship among wildtype adult males shown by *D. santomea* in naturalistic settings is, to the best of our knowledge, novel in the *Drosophila* genus. Importantly, *D. santomea* male-directed courtship is not due to sex misidentification, because individual flies that courted and sometimes also attacked males immediately switched to exclusive courtship with significantly increased vigor when encountering a female (Fig. 1K-M). In a trio assay for target sex preference, *D. santomea* males also exhibited a slight preference for courting females over males (Fig. 1N).

### The abundant and monomorphic cuticular pheromone 7-tricosene is sexually attractive to *D. santomea* males

Five of the nine *melanogaster* subgroup species (*D. simulans*, *D. santomea*, *D. yakuba*, *D. teissieri*, and *D. orena*) are considered pheromonally monomorphic because the same cuticular hydrocarbon (CHC) is the most abundant compound on the cuticles of females and males (*65–67*). For all five the defining monomorphism is of (Z)-7-tricosene (7-T) (Fig. 2A, fig. S6), a nonvolatile compound detected via contact chemosensation (*68*, *69*). *D. santomea* courts in the dark (although copulation rate is reduced) (Fig. 2B) (*70*), and removing CHCs in bulk from either females or males strongly diminishes their attractiveness (Fig. 2C), implying the existence of courtship-promoting pheromone(s) shared between sexes. That 7-T is an aphrodisiac signal for *D. santomea* males is supported by monomolecular add-back following bulk CHC removal from *D. santomea* females (Fig. 2D), ectopic addition to intact females of species lacking endogenous 7-T (*D. melanogaster* and others) (Fig. 2E, fig. S7), and positive correlation between 7-T presence and courtship elicited from *D. santomea* males in a variety of heterospecific male-male pairings (Fig. 2F). Thus, 7-T appears necessary and is sufficient to evoke sexual attraction in *D. santomea* males. The inversion from 7-T suppressing courtship in *D. melanogaster* (where it is male-specific) (*71–74*) to promoting courtship in *D. santomea* is shared with *D. yakuba* (*32*), suggesting that the 7-T valence switch arose in tandem with or soon after monomorphism in the *D. santomea / D. yakuba* common ancestor on mainland Africa (Fig. 2G) (*75*).

**Figure 2.**
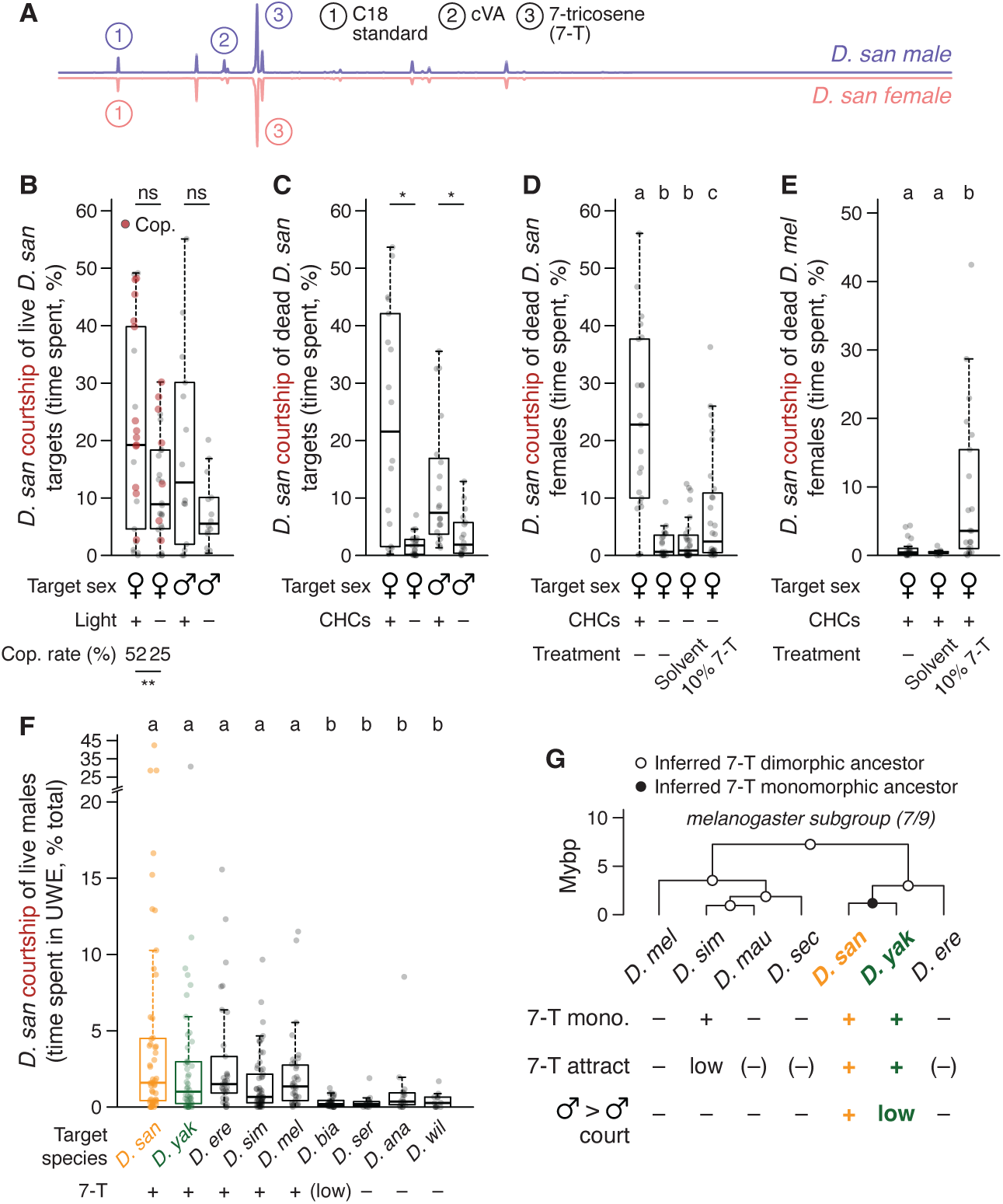
The abundant and monomorphic cuticular pheromone 7-tricosene is sexually attractive to *D. santomea* males. **(A)** Mirrored gas chromatograms representing male (purple, top) and female (pink, bottom) *D. santomea* cuticular hydrocarbon (CHC) profiles. Individual measurements made on hexane extracts from ten pooled, age-matched adult flies (3 male, 2 female replicates). Peaks representing the internal standard (C18, 1), male-specific (Z)-11-octadecenyl acetate (cVA, 2), and monomorphic (Z)-7-tricosene (7-T, 3) are indicated. **(B)** Fraction of time *D. santomea* males spend courting conspecific females or males in ambient light vs. dark during ten-minute interactions. Boxplots show full distribution range within whiskers (excluding statistically identified outliers) and second and third quartiles within boxes, with medians in bold. Individual data points (including outliers) overlayed as gray or, for males courting females that successfully copulated, red dots. Outliers retained in all summary metrics and statistical comparisons here and throughout. Note slight but non-significant decreases in courtship toward both sexes in the dark (Mann-Whitney *U* tests), although a significantly reduced copulation rate with females is observed (binomial test). **(C)** Fraction of time *D. santomea* males spend courting conspecific dead females or males with or without endogenous CHCs (removed by hexane washes) during ten-minute interactions. Significantly reduced courtship without CHCs for both target sexes (Mann-Whitney *U* tests). **(D)** Similar to (C), with 10% 7-T add-back (20 µg/fly) or hexane solvent control to conspecific dead females after CHC removal. 7-T treatment partially restores attractiveness to *D. santomea* male testers (lettered statistical groupings assigned by post-hoc Dunn’s tests following significant Kruskal-Wallis). Cohen’s *d* effect size for 10% 7-T addition (fourth) vs. no addition after CHC wash (second), 0.692 (95% CI −0.379-0.821, p=0.401). Cohen’s *d* effect size for solvent addition (third) vs. no addition after CHC wash (second), 0.221 (95% CI 0.076-1.31, p=0.042). Cohen’s *d* effect size for 10% 7-T addition (fourth) vs. solvent addition (third), 0.617 (95% CI 0.069-1.17, p=0.039). **(E)** Fraction of time *D. santomea* males spend courting dead *D. melanogaster* females (endogenous 7-T low) perfumed with 10% 7-T or solvent control (hexane). 7-T addition elicits courtship from *D. santomea* males. Significance by Dunn’s tests. **(F)** Fraction of time *D. santomea* males spend courting conspecific (orange) or heterospecific males (*D. yakuba* green, all others black) during twenty-minute interactions. 7-T presence in each species from GC-MS and phylogeny shown below, statistical groupings by Dunn’s tests above. Note correlation between 7-T presence and elevated *D. santomea* courtship. **(G)** Summarized indications of 7-T monomorphism (*65*, *66*), evidence for 7-T as a sexual attractant (*32*, *71*, *72*), and binarized male-male courtship expression in *melanogaster* subgroup species arranged by phylogeny. “(–)” indicates dimorphic species for which 7-T is not expected to be sexually attractive but has not yet been tested. Pheromone status of common ancestors at phylogeny branchpoints inferred by parsimony.

### Selectively low cVA abundance on *D. santomea* males amplifies intermale courtship

If *D. santomea* sexual attraction uses an abundant and monomorphic signal (7-T), then it would seem unsurprising that male-directed courtship is a corollary outcome of this relatively simple pheromone arrangement. However, intermale courtship in *D. yakuba* and other 7-T monomorphic species is rare (Fig. 2G) (*30*, *32*, *76*), hinting that *D. santomea* males have either gained an additional courtship-promoting signal absent in the other species or lost a courtship-inhibiting signal otherwise present. Consistent with the latter, we found that *D. santomea* males produce significantly less (Z)-11-octadecenyl acetate (cVA) than other subgroup members (84-92% reduction, 89% less than *D. yakuba*) (Fig. 3A, fig. S8). cVA is a semi-volatile olfactory pheromone produced in the ejaculatory bulb of male drosophilids (*67*, *77*, *78*) and has pleiotropic social roles including promoting aggregation (*79–81*) and aggression (*82–85*) and suppressing courtship (*86–90*), either natively on males or after transfer to females during copulation. Behavioral dose-response curves with cVA applied onto a piece of filter paper indicated that cVA indeed suppresses courtship in *D. santomea* (Fig. 3B-D), with the primary effect being to increase latency to initiation (Fig. 3C). In trios, *D. santomea* tester males showed nearly universal preference for courting solvent control treated males or females over flies directly coated with cVA (Fig. 3E). Triggering fictive cVA sensation in *D. santomea* males by phasic optogenetic activation of Or67d-expressing olfactory sensory neurons (OSNs) (*89*, *91–95*) was sufficient to interrupt natural courtship towards conspecific partners of both sexes (Fig. 3F-I, fig. S9), as also observed in *D. melanogaster* (*96*). Interestingly, the photostimulation light intensity needed to pause ongoing courtship was lower with male targets than with females (Fig. 3J), consistent with some courtship suppression even by the low endogenous amount of cVA present on *D. santomea* males.

**Figure 3.**
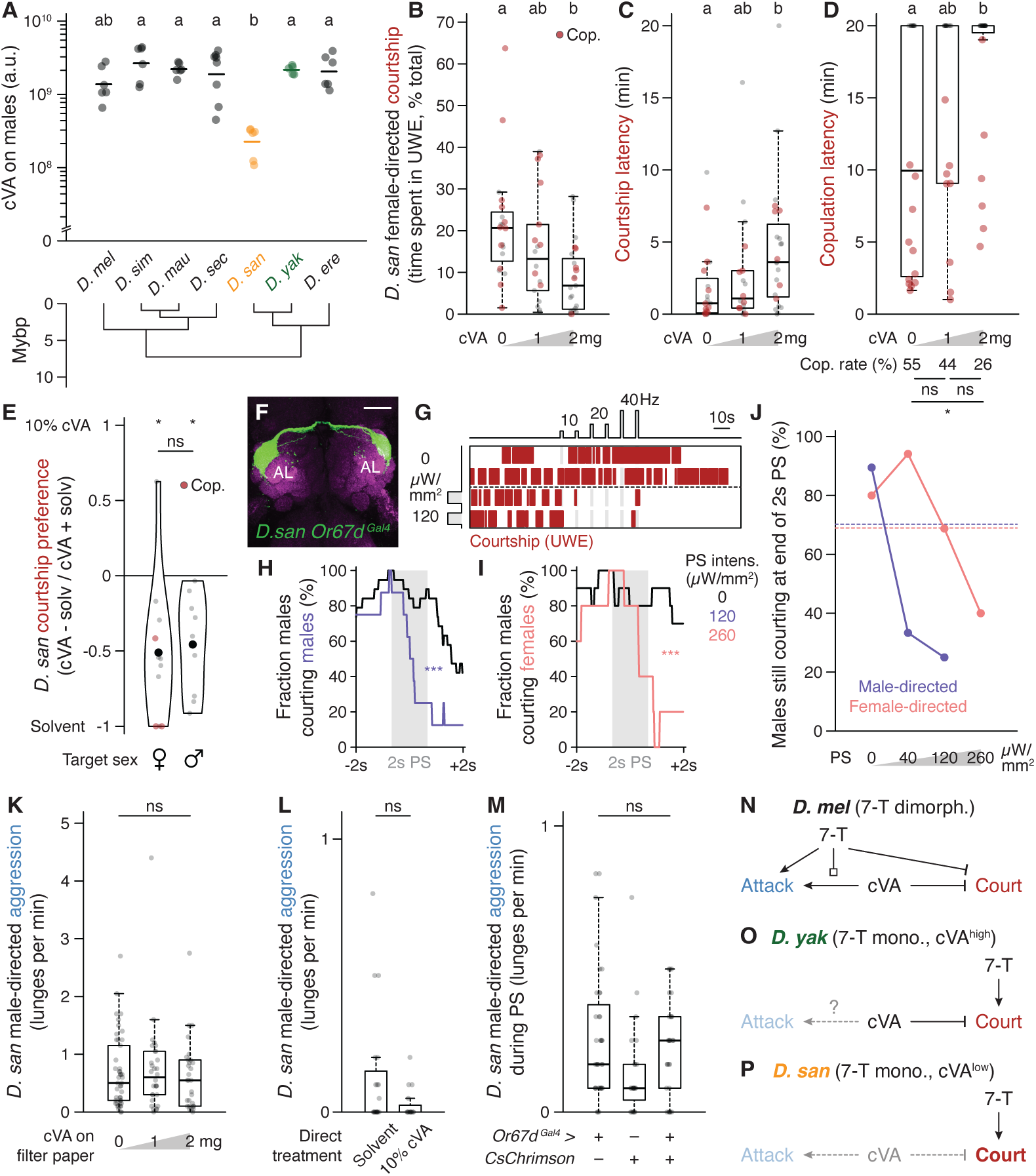
Selectively low cVA abundance on *D. santomea* males amplifies intermale courtship. **(A)** cVA abundance on males of the *melanogaster* subgroup measured in single flies (5-8 per species) by TD-GC-MS. Statistical groupings by Dunn’s tests; “a.u.” arbitrary units. Note low cVA level selectively in *D. santomea* males. Data derived from (*67*) with permission from the authors. **(B-D)** *D. santomea* sexual behavior during twenty-minute interactions with females in the presence of increasing cVA on a piece of filter paper nearby (0, 1, or 2 mg). (B) Fraction of time spent courting. (C) Latency to first courtship bout. (D) Latency to copulation. Pairwise statistical significance by Dunn’s tests, binomial tests for copulation rates. Note decreased courtship, increased latency, and decreased copulation rate at the highest cVA abundance (2 mg). **(E)** Preference index for *D. santomea* courtship towards 10% cVA (20 µg/fly) vs. solvent control (acetone) perfumed live target female or male conspecific flies during twenty-minute trio interactions. Index calculated as the difference in time spent courting each target normalized by the sum. Significance between distributions and between each distribution and a null median of zero (indicating no preference) determined by two- and one-sample Mann-Whitney *U* tests, respectively. Note that *D. santomea* male testers exhibit clear aversion to cVA-perfurmed targets of either sex. All three observed copulations occurred with the solvent control female. **(F)** Photomicrograph showing labeling pattern of *Or67d^Gal4^* in the antennal lobes (AL) of a *D. santomea* male. Immunostaining for Gal4-dependent cytoplasmic tdTomato pseudocolored green with Bruchpilot synaptic counterstain (using nc82 monoclonal antibody) in magenta. Uniglomerular labeling pattern in each AL matches the expected size and position of cVA-sensitive DA1 (*145*). Scale bar, 25 µm. See table S3 for genotypes of transgenic strains used in figures throughout. **(G)** Courtship rasters (red) for a single-housed *D. santomea* male carrying *Or67d^Gal4^* and Gal4-dependent CsChrimson paired with a group-housed wildtype conspecific male target. Two representative sham trials (above dash) and two photostimulation (PS) trials (below dash) are shown. Two-second PS blocks repeat every ten seconds for a total of six activations after a one-minute baseline. PS frequencies increase in pairs from 10 to 40 Hz within each trial and increase in intensity across trials. Note that PS immediately interrupts spontaneous courtship by tester males in experimental but not sham trials. **(H,I)** Quantification of (G) showing fraction of *Or67d^Gal4^* tester males courting males (H) or females (I) during six-second windows surrounding PS blocks (H, purple; I, pink) or sham controls (black). Data filtered for blocks where tester males showed spontaneous courtship within a half second of PS onset. Data from all three frequencies were grouped since they gave similar results but separated by intensity: 120 (H) or 260 µW/mm^2^ (I). Most tester males cease courtship by PS offset in experimental trials (significant for both target sexes by Kolmogorov-Smirnov tests). **(J)** Summary of courtship suppression effects in *Or67d^Gal4^* tester males as a function of PS intensity with male (purple) and female (pink) targets. Baselines (horizontal dashed lines) calculated as the fraction of spontaneous courtship bouts in sham trials lasting more than two seconds. Note strong effects with male targets at low intensities but only at the strongest intensity with females. **(K)** *D. santomea* male aggression during twenty-minute interactions between single-housed males in the presence of increasing cVA on filter paper. Not significant by Kruskal-Wallis test. **(L)** *D. santomea* male aggression towards males directly perfumed with 10% cVA or solvent control (acetone) during twenty-minute trio interactions. Not significant by Mann-Whitney *U* test. **(M)** Aggression in *D. santomea* males carrying *Or67d^Gal4^*and Gal4-dependent CsChrimson or genetic controls during twelve-minute photoactivation trials. Pairs of group-housed males are exposed to six thirty-second photostimulation (PS) blocks with fixed intensity and monotonically increasing frequency (1-40 Hz), with a two-minute baseline and one-minute inter-block intervals. Measurements represent cumulative lunge counts from all six PS blocks. Note insufficiency of *Or67d^Gal4^* activation to induce attack (no significance between genotypes by Kruskal-Wallis test). **(N-P)** Logic models for control of male social behaviors by 7-T and cVA in *D. melanogaster* (7-T dimorphic, N), *D. yakuba* (7-T monomorphic, O), and *D. santomea* (7-T monomorphic with reduced cVA, P). In *D. melanogaster* where 7-T inhibits intermale courtship, 7-T also gates the effect of cVA to promote aggression (*72*). A switch in 7-T function from suppressing to stimulating courtship in monomorphic species is predicted to reduce or eliminate the aggression-promoting effect of cVA without impacting its courtship-suppressing effect, as evident in *D. santomea.* Intermale courtship promoted by 7-T in *D. santomea* is further amplified due to reduced inhibition by cVA. Edge terminating in white box indicates a permissive “gating” effect by 7-T on cVA-induced aggression, in addition to its direct effect.

Unlike its courtship-inhibiting function, cVA’s aggression-promoting function in *D. melanogaster* (*82*, *83*) does not appear to be conserved in *D. santomea* males. cVA addition experiments, either onto filter paper or directly onto live target males, showed no effect of inducing aggression (Fig. 3K,L) nor did strong and direct photoactivation of Or67d OSNs (Fig. 3M). Pairings with unmanipulated heterospecific male partners presenting high endogenous cVA levels also failed to significantly increase attack behavior by *D. santomea* males (fig. S10). The ineffectiveness of cVA to promote aggression in *D. santomea* males may be due to the 7-T valence switch described above (Fig. 3N-P) or could reflect a reduced behavioral sensitivity to the pheromone relative to *D. melanogaster*.

The sexual attraction promoted by monomorphic 7-T and low levels of the male-specific anti-aphrodisiac cVA thus provide a pheromonal basis for selectively elevated courtship among *D. santomea* males. One curious additional effect was observed, presumably also due at least in part to the low cVA abundance on *D. santomea* males. When paired with males of other *melanogaster* subgroup species that find cVA sexually aversive (*32*, *67*, *89*) and show minimal sexual behavior in conspecific male-male pairs (*D. melanogaster*, *D. simulans*, *D. yakuba*), *D. santomea* males elicited 200-580% higher levels of courtship and 53-91% reduced levels of aggression from the heterospecific males relative to the amount those species showed in conspecific settings (fig. S11). Thus, *D. santomea* male attractiveness extends even across species boundaries to multiple other drosophilids.

### Central aggression circuits in *D. santomea* can evoke intense attack when photoactivated

*D. santomea* intermale sexual behavior is frequent but does not fully saturate the time that two males interact (Fig. 1B). Even so very little attack occurs outside of courtship periods. This observation, in addition to the ineffectiveness of cVA to promote aggression, prompts the question of whether *D. santomea* aggression circuitry is conserved and functional.

Multiple aggression-promoting neurons have been identified in *D. melanogaster* males (*97*, *98*), among which are P1/pC1 (and subsets therein) (*57*, *85*, *96*, *99–105*) and AIP (also called CL062) (*106–108*). We generated transgenic reagents necessary to access P1/pC1 (referred to here as P1^dsx^ to reflect the *71G01^DBD^;dsx^AD^*split driver combination used) and AIP in *D. santomea* by combination CRISPR-Cas9 knock-in (*109*, *110*) and phiC31 recombination into existing landing site lines (*111*, *112*), and tested photoactivation effects in single or paired flies using CsChrimson as the opsin effector (*113*). In comparison to *D. melanogaster* (*106*), AIP neurons had the expected position and morphology in *D. santomea* males (Fig. 4A), and photoactivation in solitary flies elicited bilateral wing elevation (i.e. threat display, the major non-contact-mediated form of aggression in *Drosophila*) (*41*, *49*, *50*) time-locked to the onset and offset of photostimulation (Fig. 4B-D, fig. S12). P1^dsx^ neurons were similarly present with a morphology resembling that in *D. melanogaster* (Fig. 4E) (*114*, *115*) and photostimulation evoked reliable time-locked courtship measured by unilateral wing extension in solitary *D. santomea* males (Fig. 4F, fig. S13).

**Figure 4.**
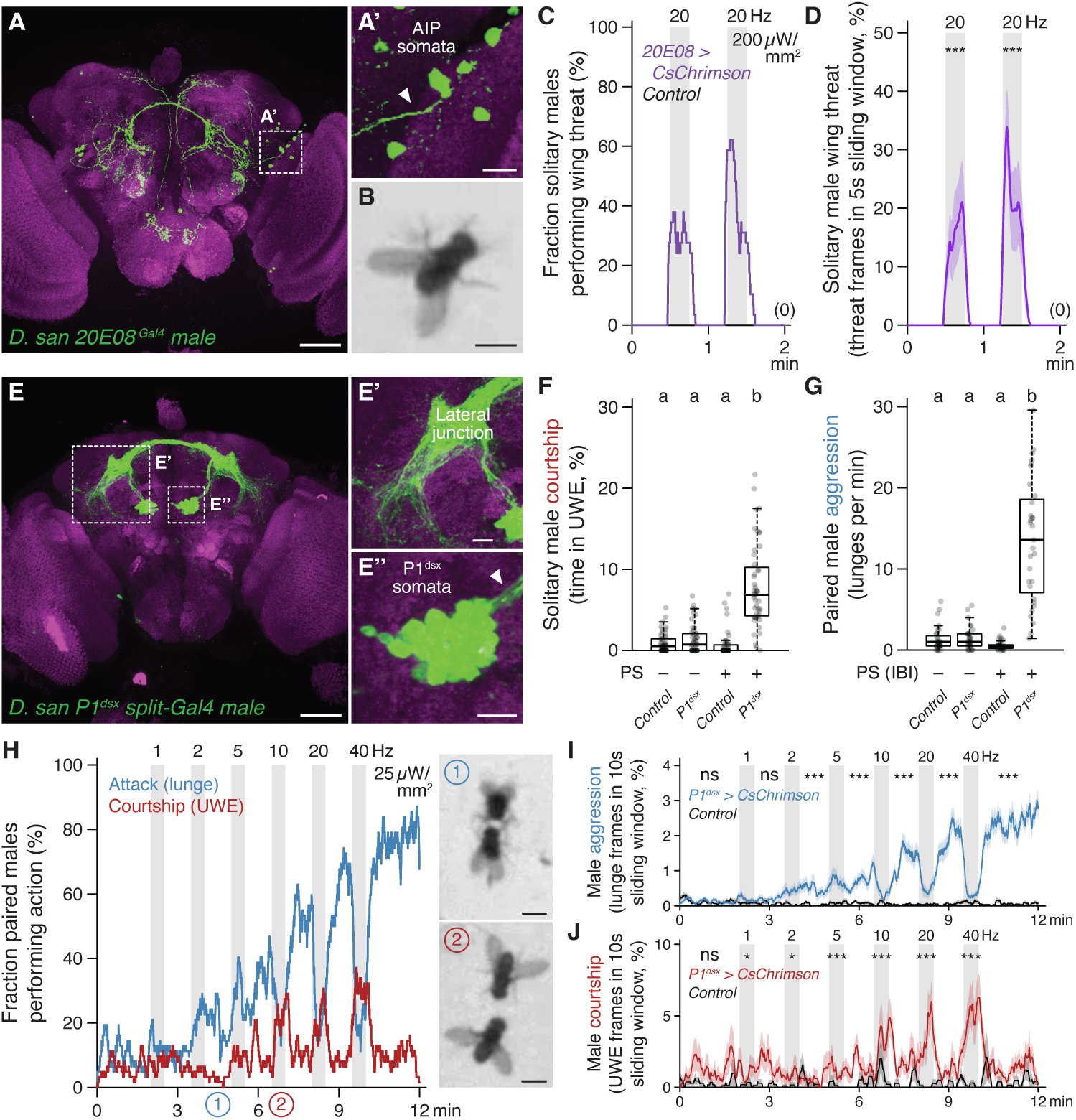
Central aggression circuits in *D. santomea* can evoke intense attack when photoactivated. **(A)** Photomicrograph showing labeling pattern of *20E08^Gal4^* in the central brain of a *D. santomea* male. Immunostaining for Gal4-dependent cytoplasmic tdTomato (green) with synaptic counterstain (nc82, magenta). Close-up (A’) shows AIP somata and axons (arrowhead) from boxed region. Scale bars, 50 µm (A), 10 µm (A’). **(B)** Video still of bilateral wing elevation (threat) elicited by PS in a solitary *D. santomea* male carrying *20E08^Gal4^* and Gal4-dependent CsChrimson. Scale bar, 1 mm. **(C,D)** Quantification of PS effects in solitary *20E08^Gal4^* tester males (purple) and genetic controls (black) exposed to two fifteen-second PS blocks with thirty-second baseline and inter-block interval. (C) Fraction of flies showing threat (penetrance). (D) Mean fraction of frames with flies showing threat in a five-second sliding window with s.e.m. envelopes (expressivity). Significant induction of threat during PS in *20E08^Gal4^* by Mann-Whitney *U* tests. **(E)** Photomicrograph showing *P1^dsx^* split-Gal4 (*71G01^DBD^;dsx^AD^*) labeling in the central brain of a *D. santomea* male. Immunostaining for Gal4-dependent cytoplasmic tdTomato (green) with synaptic counterstain (nc82, magenta). Close-ups show a characteristic region of *P1^dsx^* neuropil (lateral junction, E’) and somata and axons (E’’) from boxed regions. Scale bars, 50 µm (F), 10 µm (F’,F’’). **(F,G)** Courtship elicited in solitary (F) and aggression in paired (G) *D. santomea* males carrying *P1^dsx^* split-Gal4 and Gal4-dependent CsChrimson or a genetic control during twelve-minute photoactivation trials. In both cases group-housed males are exposed to six thirty-second PS blocks with fixed intensity and monotonically increasing frequency (1-40 Hz), with a two-minute baseline and one-minute inter-block intervals (as in H). “PS–” represents the two-minute baseline period prior to first PS. “PS+” cumulatively represents all six PS blocks (for courtship, G) or all five inter-block intervals (IBIs) and period following final PS (for aggression, F). See behavior dynamics in (H) for justification of aggression quantification during IBIs. Note significant PS induction for both actions (Dunn’s tests). **(H)** Fraction of flies showing attack (lunge, blue) and courtship (UWE, red) in same-genotype pairs of *P1^dsx^* males during six PS blocks with fixed intensity and increasing frequencies. Note highly penetrant attack mostly during light-off periods punctuated by time-locked courtship during high frequency PS blocks. Video stills at right show aggressive tussling following 2 Hz PS (1) and simultaneous courtship by both tester males during 10 Hz PS (2). Scale bars, 1 mm. **(I,J)** Expressivity and temporal dynamics of attack (I) and courtship (J) elicited by PS in *P1^dsx^* male pairs (I, blue; J, red) or a genetic control (black). Significant attack induction during inter-block intervals and after final PS (I) or courtship during PS compared to control (J) by Mann-Whitney *U* tests.

In same-genotype pairs of *D. santomea* males both experiencing titrated photoactivation, P1^dsx^ stimulation induced a mix of aggressive and courtship actions similar to the effects seen in *D. melanogaster* (Fig. 4G-J, fig. S14) (*57*). Lunge attacks were detected during low frequency stimulations (significantly at 2 Hz) and escalated in intensity as the stimulus frequency increased (Fig. 4H,I, fig. S14A-C). Fighting remained persistently elevated following the final photostimulation period for the duration of the recording. Thirty-second high frequency stimulations (>10 Hz) at one-minute intervals interrupted the otherwise monotonic increase in attack intensity with temporary phases of courtship and locomotor arrest time-locked to photostimulation (Fig. 4H,J, fig. S14D,E). Mixed-genotype pairs (in which one of the two photoactivated males was replaced with a wildtype male prepared to be passive during the assay by group-housing) confirmed that stimulated aggression was fly-autonomous and not exclusively caused by social feedback (fig. S15). However, these experiments also revealed some contextual flexibility in the expression of optogenetically evoked attack. P1^dsx^ photoactivated flies in mixed pairs required slightly higher stimulation frequency to initiate attack (5 instead of 2 Hz, fig. S15A vs. Fig. 4I) and their attack intensity remained ∼50-60% lower than in same-genotype pairs throughout the trials (fig. S15B). Interestingly, increased attack latency by P1^dsx^ photoactivated males in mixed pairs did not also obtain for courtship (fig. S15C), resulting in a shift from nearly equal balance between flies exhibiting attack or courtship first in same-genotype pairs to a majority of cases where courtship preceded attack in mixed pairs (fig. S15E). This suggests that passive male opponents can, to a limited degree, favor expression of courtship instead of attack by *D. santomea* males.

Major nodes within at least two distinct aggression-promoting circuits (P1^dsx^ and AIP) are therefore conserved in *D. santomea* and both contact- and non-contact-mediated agonistic behaviors can be efficiently evoked by their photoactivation. This rules out that the paucity of spontaneous attack in *D. santomea* male pairs under naturalistic conditions derives from an evolved loss of function of (at least these) aggression circuits. P1^dsx^ experiments reveal that *D. santomea* aggression can be elevated to supernormal intensity and exhibit persistence under optimal photoactivation conditions (∼23-fold higher lunge rate observed after the final photostimulation compared to the average during spontaneous interactions, Fig. 4I vs. Fig. 1C), but the direct effect of courtship dominates and strongly suppresses attack during P1^dsx^ photostimulation, as also in *D. melanogaster* (*57*). An opponent whose pheromone bouquet activates *D. santomea* P1^dsx^, especially a non-aggressive one, would therefore be expected to elicit courtship instead of attack as long as the activation perdures.

### A switch from cVA sexual attraction to aversion in *D. santomea* females

*D. santomea* has an interesting natural history (Fig. 5A-C) (*116–118*). They are endemic to the equatorial West African island of São Tomé, located 280 km from Gabon in the Gulf of Guinea. *D. yakuba* also inhabits São Tomé, and the two sibling species largely segregate along the elevation gradient from the volcanic island’s warm, dry, and open perimeter (*D. yakuba’s* range) to its cool, humid, forested summit (*D. santomea’s*). A sympatric zone is formed at their mid-elevation boundary, where both species and rare F1 hybrids can be collected that adhere to Haldane’s rule for sterility in the heterogametic sex (i.e., males) (*119*). Since *D. santomea* and *D. yakuba* males court females of both species promiscuously (*120*, *121*), male offspring sterility strongly pressures females to discriminate against heterospecific matings and maintain reproductive isolation (*122*). In flies this challenge would normally be expected to be met by females detecting species-specific acoustic features of male courtship song (*123*), but although *D. santomea* and *D. yakuba* songs are distinct (*61*, *63*), *D. santomea* females are still able to distinguish males of each species even if they are muted by wing amputation (*62*). These observations implicate additional species-identifying cues.

**Figure 5.**
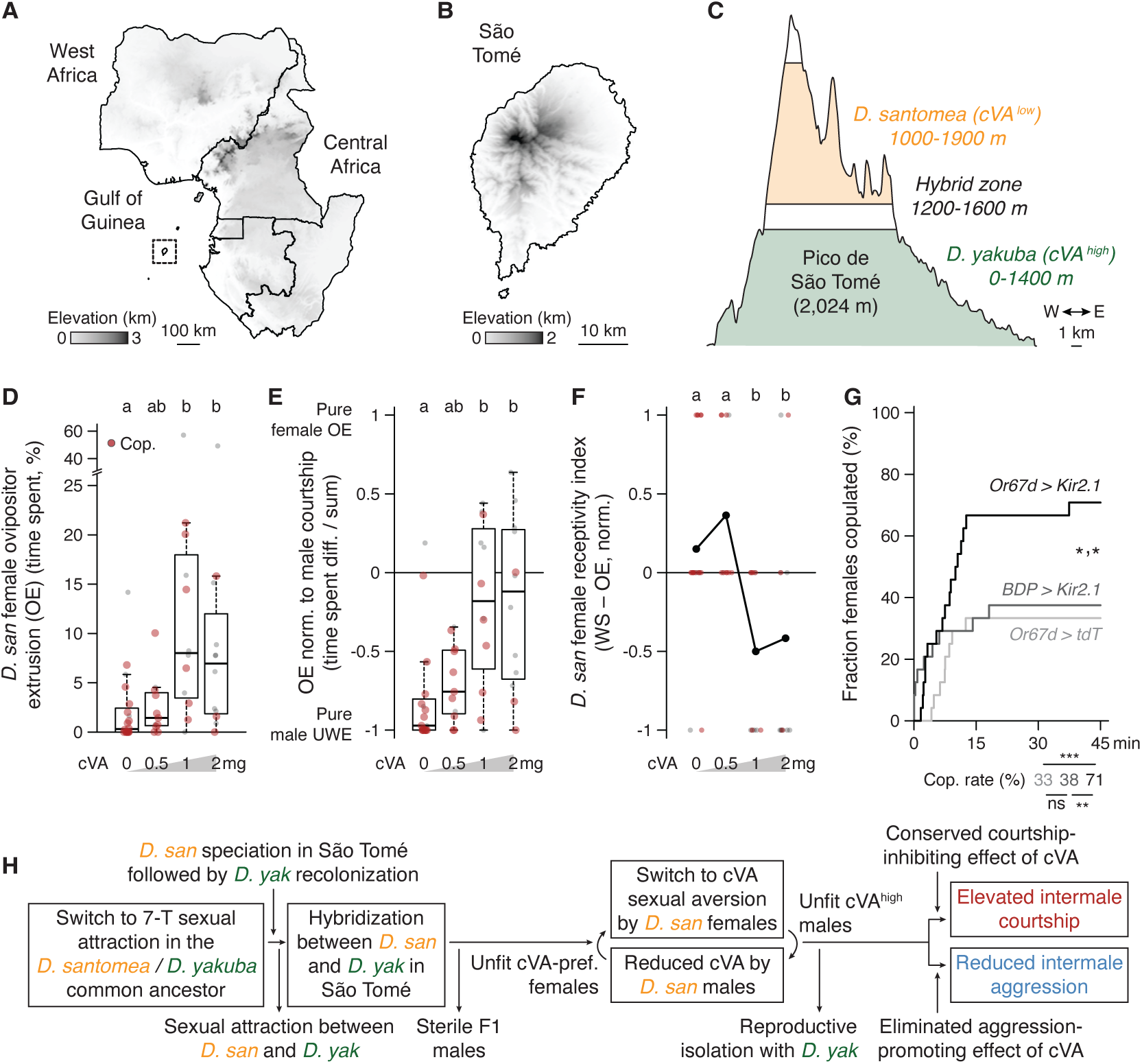
A switch from cVA sexual attraction to aversion in *D. santomea* females. **(A)** Elevation map of equatorial Central and West Africa with the volcanic island of São Tomé in the Gulf of Guinea boxed. Data from (*156*) and Terrain Tiles (Mapzen, accessed Aug. 24, 2025). **(B)** Elevation map of São Tomé. Note the central summit of Pico de São Tomé. **(C)** Latitudinal elevation profile of Pico de São Tomé showing *D. santomea* (orange) and *D. yakuba* zones (green) separated by a hybrid zone where both pure species and F1 hybrid males are found (*118*). **(D,E)** Behavioral dose-response curves of *D. santomea* female ovipositor extrusion (OE) to increasing cVA on filter paper during conspecific male courtship. (D) Female OE as fraction of total observation time. (E) Female OE normalized to male courtship (difference divided by sum of time spent by female in OE and time spent by male in UWE). Red dots, females that copulated. The lowest cVA dose induces an intermediate effect while both higher doses induce significant increases in OE expression (Dunn’s tests). **(F)** *D. santomea* female “receptivity index” as function of increasing cVA. For each female an index value of 1, 0, or −1 is possible. Observations of female wing spreading (WS, an acceptance behavior) and courtship-normalized OE (rejection, from E) were each binarized as present or absent (WS>0, OE(norm)>-0.25) and overall index per dose is calculated as the difference between the two binary values (WS-OE) normalized by the number of females scored. Note significantly decreased sexual acceptance again at the two higher cVA doses (Dunn’s tests). **(G)** Cumulative copulation latencies and rates by *D. santomea* tester females carrying *Or67d^Gal4^* and Gal4-dependent Kir2.1 (black) or genetic controls (grays) grouped with an equal number of wildtype conspecific males for 45 minutes in large chambers. “BDP” refers to a Gal4 driver without labeling evident in the central brain. 24 total females tested of each genotype (two trials each of 12 single-housed males x 12 group-housed females), significance by Kolmogorov-Smirnov tests comparing latencies or binomial tests comparing rates. **(H)** Evolutionary model for the emergence of high intermale courtship and low intermale aggression in *D. santomea,* incorporating three key pheromone changes inferred to occur prior to or following *D. santomea* speciation in São Tomé. A common sexual attractant for *D. santomea* and *D. yakuba* males (7-T) generates the behavioral potential for unfit hybridization in sympatry due to F1 hybrid male sterility (*116*). *D. santomea* females evolve sexual aversion to cVA and *D. santomea* males reduce cVA, thereby maintaining efficient intraspecific mating while also maintaining reproductive isolation from *D. yakuba* (the two events cannot be definitively ordered). Reduced male cVA, together with monomorphic 7-T, releases high intermale courtship. Reduced efficacy of cVA to promote aggression further suppresses fighting. Placement of *D. yakuba* secondary colonization after allopatric *D. santomea* speciation (instead of speciation in sympatry) from (*117*).

The cVA addition experiments above demonstrated that *D. santomea* males show increased copulation latency and reduced copulation rates in the presence of high cVA (Fig. 3C,D), but we noticed that the effects may not derive exclusively from alterations in the males’ behavior. *D. santomea* females exhibited reduced innate attraction to cVA in a place preference assay compared to *D. melanogaster* (fig. S16), and cVA exposure with courting males dramatically increased expression of ovipositor extrusion (Fig. 5D, fig. S17), a sexual rejection behavior typically seen in recently mated, unreceptive *Drosophila* females (*124–127*). Significant rejection occurred at a lower cVA dose than was required to reduce male courtship (1 vs. 2 mg, Fig. 5D vs. Fig. 3B) even after normalization to the amount of male courtship observed (Fig. 5E). Incorporating observations of wing spreading, a distinct female behavior signaling sexual acceptance (*128*), into a combined index of *D. santomea* receptivity also captured the cVA-dependent rejection response (Fig. 5F). Strikingly, Or67d OSN silencing via ectopic expression of the inward rectifying potassium channel Kir2.1 (*129*) in *D. santomea* females paired with wildtype conspecific males resulted in ∼2-fold increased copulation rates compared to controls (Fig. 5G). Sensation of endogenous cVA on *D. santomea* males therefore normally limits *D. santomea* female sexual responses.

Thus, the effect of cVA to promote sexual receptivity in *D. melanogaster* females (*89*, *130*, *131*) is reversed in *D. santomea*. This unexpected switch represents a third pheromonal change accrued in *D. santomea*, along with attraction to 7-T and reduced cVA abundance in males, relative to *D. melanogaster* and other *melanogaster* subgroup species. We hypothesize that the cVA valence reversal may help *D. santomea* females identify and avoid unwanted hybridization with high cVA-expressing *D. yakuba* males, thereby maintaining reproductive isolation in São Tomé (Fig. 5H). A question arises then of whether the increased intermale courtship that results from cVA reduction is simply a costly but necessary accommodation by *D. santomea* males to the changing olfactory preferences of conspecific females. According to this view, same-sex sexual behavior (SSB) is inherently “non-reproductive” and may be detrimental to fitness or, at best, a waste of energy (*64*). We next explored whether male-male courtship might have any adaptive value in *D. santomea*.

### P1^dsx^ activation or natural male-directed courtship induces a competitive mating advantage in *D. santomea* males

In the original comparative screen and multiple experiments conducted thereafter, paired *D. santomea* males were derived from the same stock and prepared for dyadic behavior assays identically, but nevertheless exhibited a surprising degree of courtship bias between members of the pair (Fig. 6A, fig. S18). That is, it was usually the case that just one male performed most of the total courtship observed, similar to biases observed during intermale aggression in *D. melanogaster* (*43*, *46*, *132*). In pairs where fighting also took place the two actions were most often attributable to the same fly (Fig. 6B), hinting that biased courtship and attack might both contribute to agonistic social dominance in *D. santomea*.

**Figure 6.**
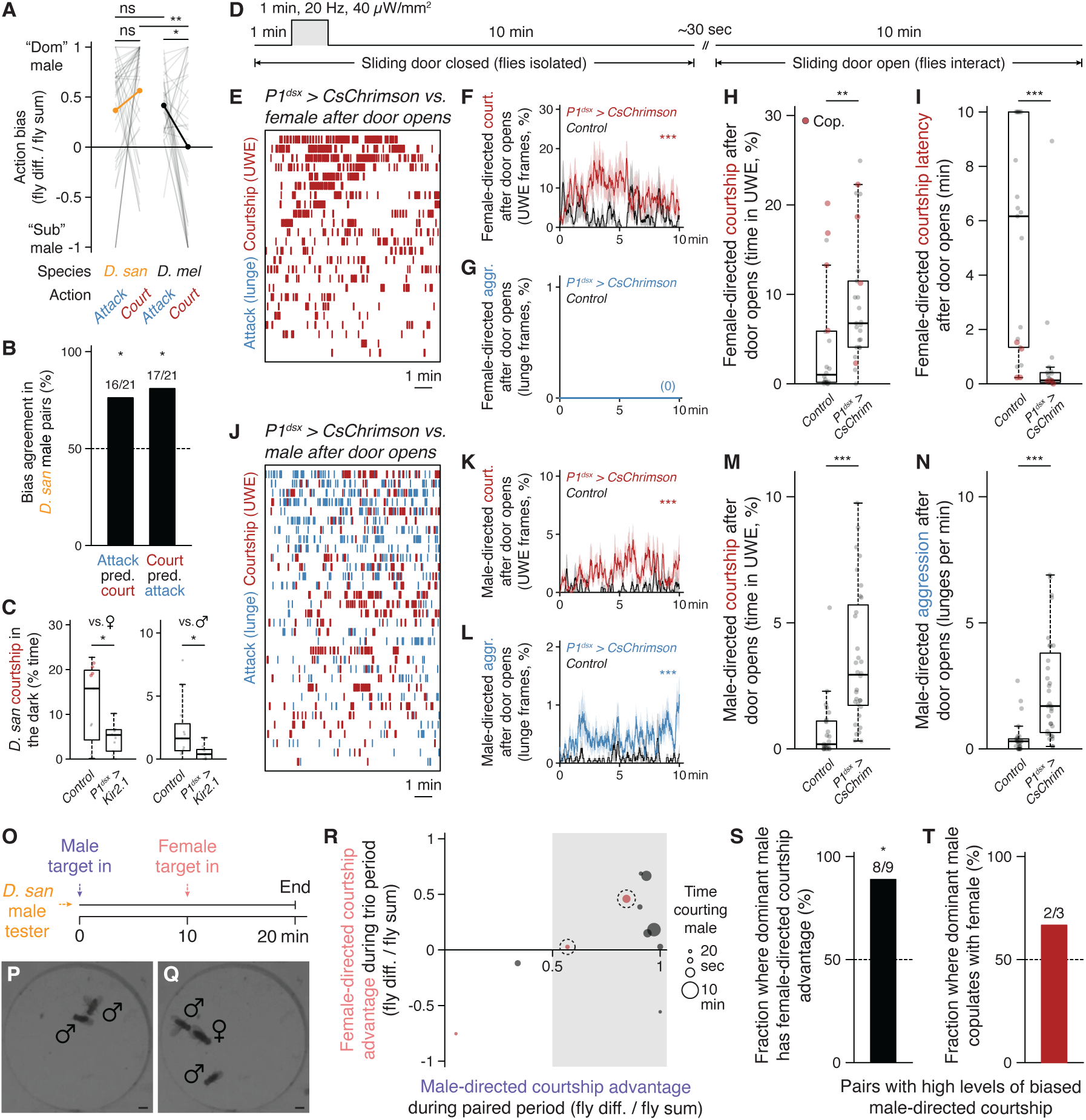
P1^dsx^ activation or natural male-directed courtship induces a competitive mating advantage in *D. santomea* males. **(A)** Spontaneous bias observed between flies in male-male *D. santomea* and *D. melanogaster* pairs for attack and courtship. Indices derived from the same fly pair are connected as lines and distribution means shown as paired orange (*D. santomea)* or black dots (*D. melanogaster*). Note high courtship bias in *D. santomea*. **(B)** Fraction of *D. santomea* male-male pairs with high attack bias where the same fly also showed majority of courtship (left), or with high courtship bias where the same fly also showed majority of attack (right). Significance compared to random chance (50%) by binomial tests. **(C)** Courtship in the dark by single-housed *D. santomea* males carrying *P1^dsx^* split-Gal4 and Gal4-dependent Kir2.1 or a genetic control paired with group-housed wildtype conspecific female (left) or male targets (right). Red dots, males that copulated (only observed in controls). Significant courtship reduction in *P1^dsx^* tester males toward both sexes by Mann-Whitney *U* tests. **(D)** Scheme for assessing persistent behavioral effects of *D. santomea P1^dsx^* split-Gal4 activation. After a one-minute baseline, isolated *P1^dsx^* tester males or genetic (non-activated) controls experience one-minute PS and a ten-minute delay. Dividers (“sliding doors”) are then removed to allow paired flies to interact spontaneously for an additional ten minutes. Pairings are between a *P1^dsx^* or control tester male and either a wildtype conspecific female (E-I) or male group-housed partner (J-N). **(E)** Behavior rasters for *P1^dsx^* tester males paired with wildtype conspecific females during the ten-minute interaction phase. Red, courtship (UWE); blue, attack (lunge, none observed). **(F-I)** Expressivity and temporal dynamics of female-directed courtship (F,H,I) and aggression (G) by *P1^dsx^* tester males compared to genetic controls during the interaction phase. Note increased courtship expression (F,H) and decreased courtship latency (I) without any coincidence of attack. Significance by Kolmogorov-Smirnov (F,G) or Mann-Whitney *U* tests (H,I). **(J-N)** Behavior rasters (J), temporal dynamics (K,L), and expressivity of male-directed courtship (M) and aggression (N) by *P1^dsx^* tester males compared to genetic controls during the interaction phase. Note significant increases for both social actions. **(O)** Scheme for determining a sexual priming effect of male-directed courtship. Two single-housed wildtype *D. santomea* males are paired for ten minutes before a wildtype female is added for an additional ten minutes (trio period). Male-directed courtship dominance is measured and identity of the dominant courter ascertained in the paired period then compared to female-directed courtship dominance in the trio. **(P,Q)** Video stills of a male courting another male first in the paired period (P) then courting a female in the trio (Q). Scale bars, 1 mm. **(R)** Quantification of the sexual priming effect (O-Q). Female-directed courtship dominance in trios (y-axis) plotted as a function of male-directed courtship dominance in pairs (x-axis). Points above the x-axis represent cases where the same male exhibited male-directed courtship dominance in pairs and female-directed courtship dominance in trios. Dot size (scale on right) represents total amount of male-directed courtship in pairs and data are filtered for pairs with at least twenty seconds of courtship during this phase, no matter the bias. Red dots, males that copulated. Gray box, region reflecting cases where males exhibited with high levels of courtship dominance during the paired period. Note enrichment of points above the x-axis in cases of high male-male courtship dominance. **(S,T)** Categorization of effects on female-directed courtship dominance (S) and copulation advantage (T) from (R). (S) All but one male exhibiting male-directed courtship dominance in pairs (89%) also exhibited female-directed courtship dominance in trios (significant by binomial test). (T) Two out of three copulations observed were by the male exhibiting male-directed courtship dominance in pairs (circled red dots in R). The exception was from a “submissive” male in which courtship bias was very weak (uncircled red dot at bottom left in R).

*D. santomea* males do not appear to be territorial in the laboratory (fig. S19), but we reasoned that the dominant courting male in dyadic male pairs might be afforded a courtship advantage in subsequent female encounters, due to persistent social arousal. Given that activity in homologous P1^dsx^ neurons is required for both female- and male-directed courtship in *D. santomea* (Fig. 6C), we first confirmed that the long-lasting, behaviorally latent internal state that is evoked by transient activation of P1/pC1 neurons in *D. melanogaster* (*57*, *97*, *100*) is also obtained in *D. santomea* (fig. S20A). Persistent effects of transient P1^dsx^ photoactivation on later social behavior were assessed using a “sliding door” assay as described previously (*57*). *D. santomea* P1^dsx^ tester or genetic control males were loaded into behavior chambers on the opposite side of a removable barrier from a wildtype target conspecific female or male prepared to be passive by group-housing. A one-minute P1^dsx^ photostimulation was performed during the initial isolation phase, during which time tester males exhibited stimulus-locked unilateral wing extension (fig. S20B-E). Barriers were removed ten minutes following stimulation offset, after which tester and target flies were allowed to interact for an additional ten minutes (Fig. 6D). P1^dsx^ tester males showed vigorous courtship, but not aggression, upon encountering target females, exceeding that observed in genetic controls and with greatly reduced latency to initiation (Fig. 6E-I, fig. S20F). When encountering a target male, P1^dsx^ tester males exhibited a mixture of courtship and aggression at levels that both also exceeded genetic controls (Fig. 6J-N, fig. S20G). Thus, brief P1^dsx^ photoactivation induces a persistent internal state that can enhance subsequent social interactions for at least a period of multiple minutes in *D. santomea*, resulting in courtship to both sexes but aggression exclusively to males. *D. santomea* males can therefore discriminate conspecific sexes and adjust social behaviors accordingly even when highly aroused.

Next, we investigated whether a naturalistic male-male social interaction, rather than P1^dsx^ photoactivation, would induce a persistent arousal state transferrable to subsequent male-female interactions. Wildtype tester males were exposed to two consecutive social encounters: first in a pair with another male, during which asymmetric courtship was displayed by one of the males, and second in a trio with the same two males and a newly added female (Fig. 6O-Q). Courtship bouts by both males were tracked in both periods (accounting for target sex in trios), to assess whether dominance in male-directed courtship during the first encounter would translate to dominance in female-directed courtship in the second. *D. santomea* males that dominated courtship in the paired male setting exhibited a significant courtship advantage toward females in the trio phase of the experiment, compared to their sub-dominant male partners (Fig. 6R,S). Copulations were rare (3/9 trios), but two of the three observed were by the previously dominant male (Fig. 6T) (in the third case, the male pair exhibited very weak dominance). Thus, an episode of male-directed courtship in *D. santomea* increases the likelihood of courtship with a subsequently encountered female, suggesting a carry-over effect of a state of P1^dsx^-evoked social arousal that generalizes to mating behavior with both sexes.

### Convergent social behaviors and pheromones in *D. persimilis* males

Our behavioral screen also uncovered a second species highly prone to male-directed courtship, *D. persimilis* (Fig. 1B, fig. S1A). *D. santomea* and *D. persimilis* last shared a common ancestor ∼50 million years ago and intermale courtship appears to be selectively derived in each species (Fig. 7A). *D. persimilis* and *D. santomea* both inhabit cool, high elevation, montane regions, *D. santomea* in West Africa and *D. persimilis* in North America, and both are members of a sibling species pair with which they geographically overlap (*53*). Importantly, males of *D. persimilis* and *D. pseudoobscura* (the sibling species) both court females interspecifically but females show strong discrimination against hybridization (*133–135*). Like *D. santomea, D. persimilis* male-directed courtship shares visual resemblance with the female-directed form and is most often biased to just one male within each pair (Fig. 7B-D, fig. S21A,C,D). *D. pseudoobscura* males instead show the more genus-typical social patterns of male-directed aggression and female-directed courtship (Fig. 7E-G, fig. S21B-D) (*136*). Finally, *D. persimilis* males have also converged on the two key pheromonal features of CHC monomorphism and low cVA expected to reduce or eliminate major chemical barriers to male-male courtship (Fig. 7H,I) (*67*, *137*). Similar phylogenies and natural histories therefore correlate with convergent pheromones and male-male sexual behaviors shared between these two distantly related and geographically separated *Drosophila* species.

**Figure 7.**
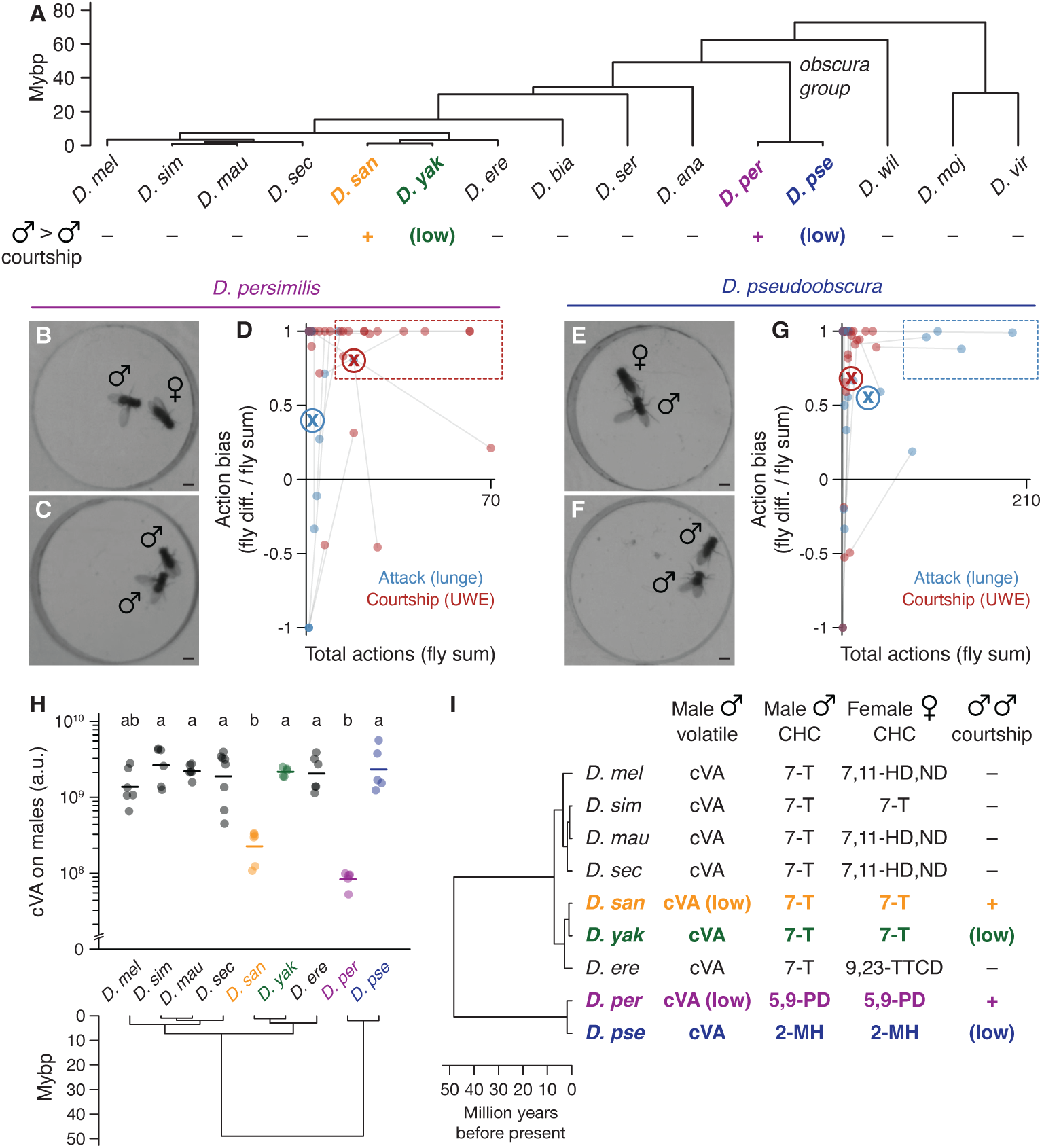
Convergent social behaviors and pheromones in *D. persimilis* males. **(A)** Summarized indications of male-male courtship presence/absence in *Drosophila* species arranged by phylogeny. Note sibling species *D. persimilis* (purple) and *D. pseudoobscura* (blue) in the obscura group. **(B,C)** Video stills of *D. persimilis* males courting conspecific female (B) and male targets (C). Images selected to represent species-typical action selection. Scale bars, 1 mm. **(D)** Action biases in *D. persimilis* male-male pairs. Male-directed courtship (red dots) is more common than attack (blue dots) and often shows strong bias to one fly (boxed region). Attack and courtship measurements from the same pair connected by thin lines. Circled “X” indicates mean value for the corresponding action. **(E-G)** Video stills (E,F) and action biases (G) for *D. pseudoobscura* as for *D. per*. Note female-directed courtship (E) and biased male-directed attack (F,G). Scale bars, 1 mm. **(H)** cVA abundance on males of the *melanogaster* subgroup and *D. persimilis* / *D. pseudoobscura* by TD-GC-MS (*67*). Statistical groupings by Dunn’s tests; “a.u.” arbitrary units. Note low cVA levels in *D. persimilis* and high in *D. pseudoobscura* as in *D. santomea* / *D. yakuba*. **(I)** Summary of relevant pheromones (*67*, *76*, *137*) and social behaviors for the *melanogaster* subgroup (excluding *D. teissieri* and *D. orena*) and the *D. persimilis* / *D. pseudoobscura* sibling species pair. CHC monomorphism is common to multiple species but low cVA abundance and high male-male courtship are shared only between *D. santomea* (orange) and *D. persimilis* (purple). Information compiled from the current study and prior sources.

## Discussion

Evolutionary changes in the expression and central processing of cuticular hydrocarbons (CHCs) have been shown to underlie phylogenetic variation in pheromonal control of male-female courtship in *Drosophila* (*22*, *67*, *138*). For example, changes in the sex-specificity of pheromone production can switch species from dimorphic to monomorphic CHC patterns (*28*, *139*), as is inferred to have occurred at least twice in the *melanogaster* subgroup (*30*, *32*). Reversal in the behavioral valence of certain CHCs can also alter patterns of sexual attraction and aversion, as for 7-tricosene (7-T), which inhibits male courtship in dimorphic *D. melanogaster* (*71–74*) but promotes courtship in monomorphic *D. yakuba* (*32*) and, to a lesser extent, *D. simulans* (*27*, *30*, *71*).

Here we have identified a set of three evolutionary changes to pheromone signaling in *D. santomea*: a switch to monomorphism and sexual attraction to 7-T in males, a roughly ten-fold reduction of the olfactory signal cVA relative to other *melanogaster* subgroup species also in males, and a behavioral valence switch to cVA in females from promoting (*89*, *130*, *131*) to inhibiting sexual receptivity. Together these changes can explain the striking and selective amplification of *D. santomea* intermale courtship and illustrate how multiple changes in the behavioral valence or abundance of pheromones in both sexes can collectively drive evolutionary variation in intrasexual social behaviors. Though definitive evolutionary causes for the three changes are difficult to resolve retrospectively, strong selection pressure on *D. santomea* to maintain reproductive isolation from *D. yakuba* in their overlapping habitat in São Tomé (West Africa) offers a unifying and plausible explanation for the emergence and fixation of these changes. A similar pattern of pheromonal and intermale behavioral changes is shown by *D. persimilis* in North America, whose females also need to maintain reproductive isolation from the sympatric sibling species *D. pseudoobscura* (*53*), implying generality and convergent evolution of responses to similar selection pressures in these two drosophilid species.

### A layered evolutionary path to elevated intermale courtship in *D. santomea*

*D. melanogaster* males (*50*, *97*) and males of most *Drosophila* species surveyed here exhibit primarily female-directed courtship and male-directed aggressive behavior. The sex-specificity of these social behaviors in dimorphic *D. melanogaster* is controlled in part by male-specific 7-T and cVA, which function similarly to suppress intermale courtship and promote aggression (*69*, *72*, *82*, *83*). However, *D. santomea* and *D. yakuba* are monomorphic for 7-T (*65–67*) and males show 7-T sexual attraction rather than aversion (*32*). This permits these species to pursue females and mate in the dark like dimorphic species can, but shifts the function of sex discrimination to other pheromones or sensory modalities. Despite high 7-T abundance and monomorphism, intermale sexual behaviors in these species are usually minimal. The rare evolutionary occurrence (within our screen, at least) of elevated intermale courtship in *D. santomea* parallels a selective reduction in the abundance of cVA, an olfactory signal which in *D. melanogaster* suppresses male courtship (*71*, *86–90*) and is otherwise present at high levels on males of *melanogaster* subgroup species. That this correlation reflects causation is demonstrated by the fact that *D. santomea* male courtship behavior can be suppressed either by addition of exogenous cVA or by direct photoactivation of olfactory sensory neurons (OSNs) expressing the cVA receptor Or67d (*89*, *91–95*). Thus, the high level of sexual behavior among *D. santomea* males can be explained by two evolutionary changes in distinct cuticular and olfactory pheromone signaling pathways: a switch in the behavioral valence of 7-T (shared with *D. yakuba*) and a reduction in the abundance of cVA (specific to *D. santomea*).

A potential evolutionary explanation for the derived male cVA reduction in *D. santomea* was lacking until our unexpected discovery of a second sexual valence switch (and third pheromonal change in total), this time in the behavioral response to cVA by females. cVA promotes mating receptivity in *D. melanogaster* females (*89*, *130*, *131*) but has the opposite effect in *D. santomea*. The switch may be related to interspecific sexual dynamics between *D. santomea* and *D. yakuba* at their geographic boundary in São Tomé. *D. yakuba* males court *D. santomea* females vigorously (*120*, *121*) and rare (sterile) F1 hybrid males can be collected within the sympatric zone (*116–118*). Thus, we speculate that the *D. santomea* female cVA valence switch arose due to ongoing selection pressure to maintain reproductive isolation from *D. yakuba* (*122*). If this explanation is correct, it would imply that *D. santomea* females use the high level of cVA on *D. yakuba* males as a sex- and species-identifying cue to ward off unwanted advances from heterospecific suitors (though we cannot exclude that *D. yakuba* courtship song is also used by *D. santomea* females for species discrimination) (*61–63*). This female aversion to cVA may have led in turn to reduced cVA production by *D. santomea* males under sexual selection pressure (*140*).

However, two other evolutionary scenarios are also possible. *D. santomea* males may have initially reduced cVA abundance due to neutral drift or to pheromonally distinguish themselves from *D. yakuba* males (*141*), which in turn may have pressured *D. santomea* females to develop cVA aversion by sexual selection operating in the reverse direction. Alternatively, *D. santomea* female cVA aversion and male cVA reduction may have evolved simultaneously, potentially in genetic linkage (*142*). In any of these scenarios, the ultimate outcome would still be an elegant pheromonal mechanism which maintains efficient intraspecific mating in *D. santomea* while minimizing unfit hybridization with *D. yakuba*. Interestingly, a similar sexual reversal from cVA promoting to inhibiting female receptivity has been reported in one other drosophilid, the agricultural pest *D. suzukii* (*143*), whose males show fully eliminated cVA production (*144*). The evolutionary circumstances and selection pressures leading to *D. suzukii* cVA changes are unclear, however, as are potential consequences for intermale social behaviors.

Mechanistic explanations for the changes to cVA signaling in *D. santomea* females remain to be elucidated. In *D. suzukii*, female cVA valence reversal correlates with dramatic size reduction of the olfactory glomerulus targeted by Or67d OSNs (DA1) (*144*), but this mechanism does not seem to be shared with *D. santomea* where DA1 size and position appear similar to closely related species (*145*). In *D. melanogaster*, cVA promotes female receptivity through activation of DA1-lvPN (*84*) and pCd neurons (*100*, *146*) in the central brain. Whether behavioral valence inversion in *D. santomea* females is reflected in a physiological inversion of the effect of cVA on these neurons, from excitatory to inhibitory, is an interesting possibility. A challenge also remains in resolving the genetic and biochemical nature of reduced cVA abundance in males and determining whether the divergent *D. santomea* and *D. persimilis* lineages achieved reductions by similar means. A biosynthetic logic for cVA production in the ejaculatory bulb has been proposed (*147*) and one elongase identified (*148*), but additional enzymes necessary to generate cVA from fatty acid precursors have not yet been identified. *Cyp6a20* has been identified as a potential degradative enzyme for cVA which is expressed by non-neuronal cells in Or67d-expressing sensillae (at1) (*44*, *47*), and may be another target of selection.

### Mechanisms for suppressing aggression in *D. santomea* males

*D. santomea* males show little spontaneous attack. However, photoactivation of aggression-promoting neural circuits in the central brain is sufficient to elicit multiple contact- or non-contact-mediated agonistic actions (e.g., threat displays, lunging, tussling) with supernormal intensity, ruling out a lack of vigor or selective atrophy of aggression circuits as explanations for the low level of spontaneous aggressive behavior. Because *D. santomea* males can discriminate conspecific sexes they must be able to detect male- and/or female-specific cues. Evidently, male-specific cues are either insufficient to activate aggression circuits, or their action is overridden by coincident male-derived courtship-promoting cues (e.g., 7-T) (*97*). Notably, as in *D. melanogaster* (*57*), *D. santomea* males only attack each other during the light-off periods of phasic P1^dsx^ photoactivation experiments. This OFF-activation pattern of aggression may reflect a post-inhibitory rebound mechanism following stimulation-locked courtship, perhaps resulting from reciprocal inhibition between courtship and aggression circuits (*97*, *99*). Apparently the effects of this rebound are long lasting, since propensity for male-directed attack remains elevated even ten minutes after P1^dsx^ neurons have been stimulated in the sliding door assay, as initially demonstrated in *D. melanogaster* (*57*). One key difference from *D. melanogaster* is apparent, however: *D. santomea* males also court the conspecific males they subsequently encounter, consistent with the high 7-T, low cVA, or other sexually attractive male features occasionally overriding rebound aggression by promoting courtship. This interpretation is supported by the fact that no aggression and instead exclusive courtship is performed towards females following barrier removal.

In addition to inhibiting male courtship, cVA and Or67d signaling can promote aggression among *D. melanogaster* males (*82–85*). If this were also the case in *D. santomea*, then cVA reduction alone could have parsimoniously explained both the elevated courtship and reduced fighting observed in *D. santomea* intermale social interactions. However we have so far been unable to verify this prediction, because various experimental efforts to elevate cVA including strong and direct photoactivation of Or67d OSNs have been ineffective at increasing *D. santomea* aggression (although effective at suppressing intermale courtship). A possible explanation for these negative results is suggested by the established hierarchical control of male-male social behavior by 7-T and cVA in *D. melanogaster* (*72*). There, cVA can promote fighting but only if 7-T signaling remains intact. In flies lacking 7-T or its receptor Gr32a (*69*), cVA has no effect to promote aggression or to inhibit the extensive male-male courtship unmasked by genetic disruption of 7-T signaling. Thus, suppression of intermale courtship by 7-T is apparently a prerequisite for the aggression-promoting effect of cVA in *D. melanogaster*. By extension, since in *D. santomea* 7-T no longer suppresses courtship, then according to this hierarchical model increasing cVA experimentally should be unable to promote fighting, as is observed. This hypothesis does not exclude the possibility that thus-far undetected evolutionary changes in aggression-promoting components of cVA central processing circuitry (*84*, *100*) may also prevent this pheromone from promoting aggression in *D. santomea* males.

### Recent speciation and intermale sexual behavior

It seems unlikely to be a coincidence that the two cases of highly expressive and penetrant male same-sex sexual behavior (SSB) discovered here in *Drosophila* are each observed in members of sympatric sibling species pairs susceptible to unfit hybridization. This correlation suggests that recent speciation may promote elevated SSB expression (or alternatively that high SSB may promote speciation), especially in the species inferred to be derived due to its more restricted geographic range. Interestingly, such a correlation was also found in a phylogenetic analysis of SSB incidence across all mammals, including the Hominidae (*149*). Our findings provide a general model that may explain at least some of these cases. Namely, where sympatry necessitates enforcement of reproductive isolation between interfertile species that produce unfit hybrid offspring (such as sterile F1 males) (*122*), mechanisms that evolve to prevent interspecific mating may draw upon sensory cues that, once altered, also modify intraspecific social behaviors in such a way as to release SSB. If the condition of sympatry is removed, or additional species-identifying cues are enlisted, or compensatory mechanisms evolve to alter the resulting pattern of intrasexual social interactions, then SSB may eventually disappear, unless it provides adaptive value.

Such an adaptive value for SSB may have accrued in *D. santomea* for either of two reasons, not mutually exclusive. Males performing natural courtship during male-male encounters (or courtship evoked by P1^dsx^ photostimulation) initiate courtship more quickly and sustain it more vigorously toward a female if she appears within a few minutes. Males could therefore benefit from this persistent social arousal or “priming” effect. Alternatively, the strong courtship bias observed in dyadic male-male interactions, and the fact that the same male that courts is also the one more likely to attack, suggests that intermale courtship could have an adaptive function in mediating social dominance (*64*). Interestingly, evidence in laboratory mice suggests that most episodes of male SSB reflect dominance mounting, a low-intensity form of aggressive behavior, rather than affiliative sexual behavior (*150*). SSB as a dominance behavior has also been proposed in other taxa (*151*, *152*).

The general relationship between SSB and aggression across species appears more complex. The survey study in mammals (*149*) found a positive correlation between SSB and lethal male violence, motivating the hypothesis that SSB may evolve to ease escalating intermale aggression. If this alternative model also pertains to *Drosophila*, then the sequence and causal role of evolutionary events identified here should be reinterpreted: rather than the key benefit of pheromonal (especially cVA) changes being to avoid cross-species hybridization, it may instead have been to effectively replace intermale aggression with courtship, a form of social interaction carrying reduced risk of harm to individuals (*64*). In other words, *D. santomea* males may have evolved a social strategy in which the ultimate functions of aggression that improve fitness (e.g., securing resources, mates, etc.) can be achieved by asymmetric intermale courtship instead, avoiding aggression’s injurious costs but maintaining its agonistic or competitive function (*153*). Resolving or harmonizing these alternatives, potentially by modeling or controlled selection experiments, should help to clarify the evolutionary origins of social behavior across the animal kingdom, perhaps including in *Homo sapiens*.

### Caveats

Our data implicating 7-T as a male aphrodisiac involves addition of exogenous pheromone to dead females. Therefore, we cannot exclude that other components of the CHC bouquet and/or non-CHC pheromones contribute to *D. santomea* male sexual attraction to live conspecific females or males, or that non-chemosensory signals also play a role (*30*). We have also so far been unable to demonstrate that 7-T sensation or physical or olfactory contact with unaltered conspecific females or males is sufficient to induce physiological activation of P1^dsx^ somata or neuropil in restrained *D. santomea* males using two-photon calcium imaging. We believe these negative imaging results likely reflect technical issues rather than biology, since both 7-T and conspecific females carrying endogenously high levels of 7-T can activate P1^dsx^ neuropil in the sibling species *D. yakuba* (*32*). Finally, our evidence that cVA inhibits *D. santomea* male courtship derives from gain-of-function experiments involving exogenous cVA addition or optogenetic activation of Or67d OSNs, all of which are vulnerable to behavioral outcomes resulting from supraphysiological pheromone exposure (whether chemical or fictive). It is therefore possible that cVA pheromonal signaling can have distinct effects on courtship or other social behaviors at lower concentrations or under weaker activation conditions.

## Acknowledgments

We thank the Anderson lab (Caltech) for help throughout the project, including experimental suggestions and conceptual advice. Liliana Chavarria, Gina Mancuso, Celine Chiu (Caltech) for administrative help. Arnold Sanchez (Caltech) for fly stock maintenance. Heather Dionne, Jeremy Hasseman, Elizabeth Kim, Katie Schretter, and Hiroshi Shiozaki (Janelia Research Campus) for constructs, reagents, and fly stocks. Rory Coleman (NYU) and Vanessa Ruta (Rockefeller) for constructs, fly stocks, information regarding unpublished results, and comments on the manuscript. Hong Yu and Garrett Kuiken (Rainbow Transgenic Flies) for embryo injections. Kathy Musial, Gary Roberson (Huntington Botanical Gardens), and David Orr (Waimea Botanical Garden) for access to fruits for host specialization experiments. Ryan York (Arcadia Science) for help with behavioral and evolutionary analyses. Peter Andolfatto (Columbia) for access to an assembled and annotated *D. santomea* reference genome. Markus Knaden (MPI Chemical Ecology) for permission to reanalyze published TD-GC-MS data.

## Funding

NIH NIDA 5R37DA031389 (DJA)

Howard Hughes Medical Institute (DJA)

Caltech Center for Evolutionary Science (YO, DJA)

Jane Coffin Childs Memorial Fund Postdoctoral Fellowship (YO)

Della Martin Foundation Postdoctoral Fellowship (YO)

## Author Contributions

Conceptualization: YO, DJA

Formal Analysis: YO, DJA

Funding Acquisition: YO, DJA

Investigation: YO, THN, HS, ELB

Methodology: ELB, YD, DLS

Resources: HS, MAK, YD, DLS, JP

Software: THN, HS

Visualization: YO, DJA

Writing – Original Draft Preparation: YO

Writing – Review & Editing: YO, THN, HS, ELB, MAK, YD, DLS, JP, DJA

## Competing Interests

The authors declare no competing interests.

## Data and Materials Availability

All data are available in the main manuscript or the supplementary materials. Raw source data, analytical code, constructs and associated maps, and fly stocks will be made publicly available in adherence with current Caltech and HHMI policies.

## Supplemental Figure Legends

**Figure S1.**
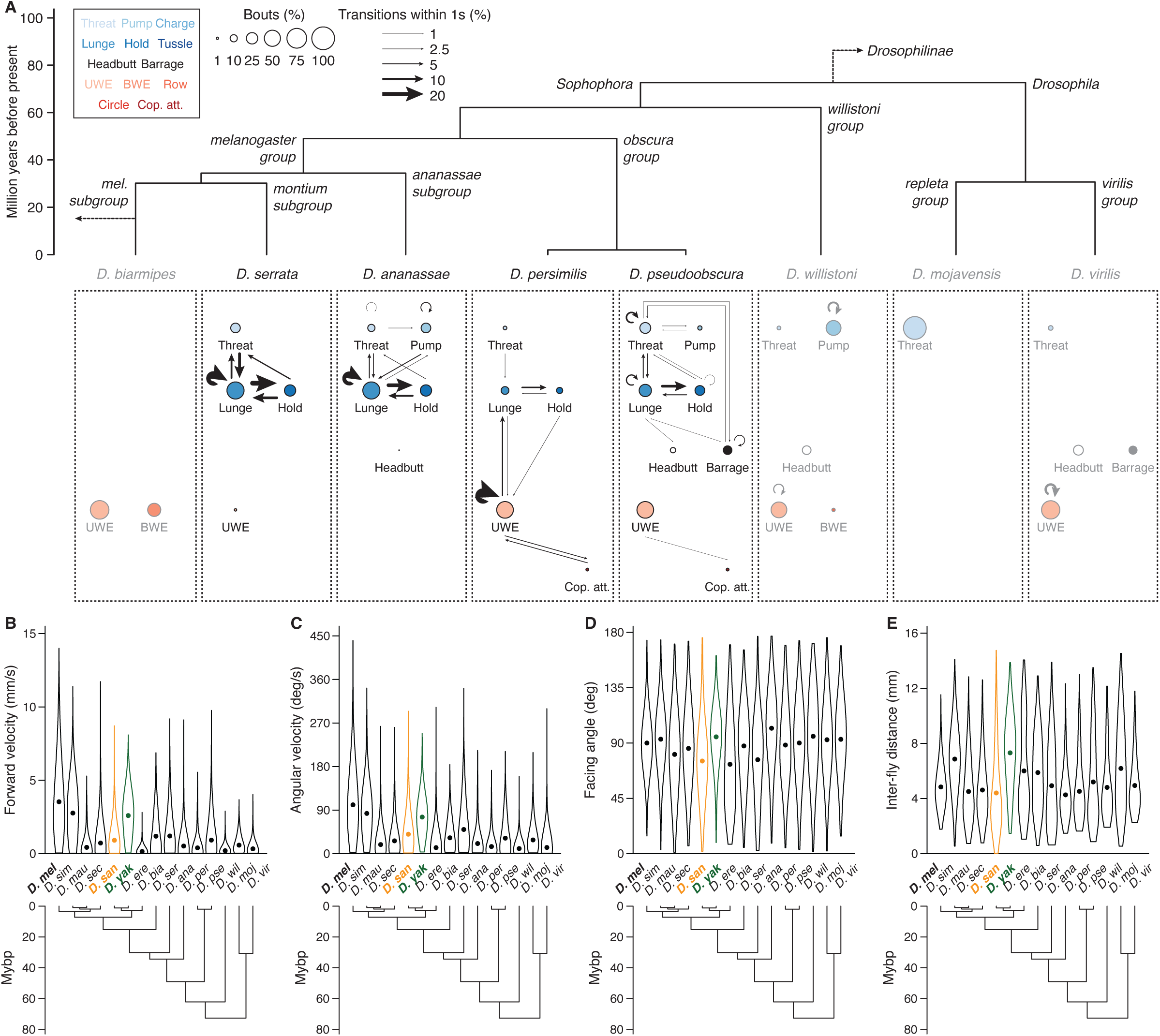
Additional behavior ethograms and locomotor analyses of *Drosophila* intermale social interactions. **(A)** Ethograms from twenty-minute interactions depicting aggressive and courtship actions scored in three representative conspecific male-male pairs per species across various *Sophophora* and *Drosophila*. Nodes are bout counts of a given action type, with sizes representing frequency normalized to summed bouts scored across all actions (expert annotation). Edges indicate action transitions, with weights representing the fraction of bouts of a given action (arrow origin) for which a second action (arrow destination) occurred within one second. Nine total actions in each ethogram are arranged in five rows: first, two aggressive threat actions (threat, pump) (*106*); second, two aggressive contact-mediated actions (lunge, hold) (*49*); third, two additional contact-mediated aggressive actions (headbutt, barrage) (*157*); fourth, two courtship actions utilizing the wings and typically directed toward females (UWE, BWE) (*20*); fifth, copulation attempt. Nodes indicating aggressive actions are filled with blue shades and courtship actions with reds, except for headbutt (white) and barrage (black). Species arrangement and divergence times according to compilation of available phylogenies (*75*, *154*, *155*). Gray coloration indicates that very few social actions of any kind were observed. Abbreviations: UWE, unilateral wing extension; BWE, bilateral wing extension; Cop. att., copulation attempt. Barrage described previously in *D. pseudoobscura* (*136*) but named here. **(B-E)** Locomotor summaries for all fifteen species included in the behavioral screen. Per-frame calculations of each fly’s forward velocity (B), angular velocity (C), facing angle relative to the partner (D), and inter-fly distance (E) are derived from automated tracking of both flies (*56*) in all recorded male-male pairs. Distributions shown as violin plots with dotted means. *D. santomea,* orange; *D. yakuba*, green.

**Figure S2.**
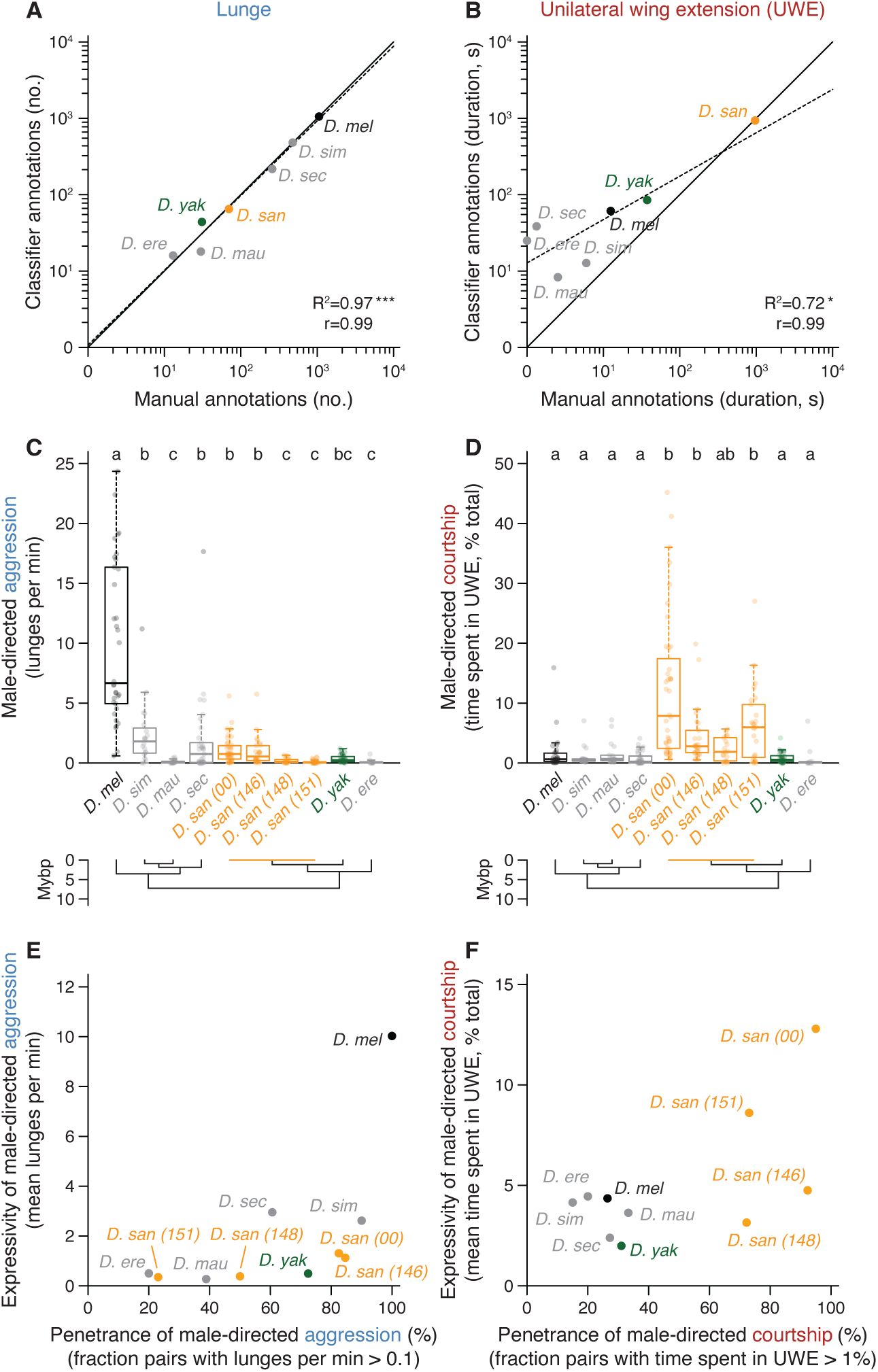
Performance and usage of automated behavior classifiers for intermale social interactions within the *melanogaster* subgroup and additional *D. santomea* strains. **(A,B)** Automated behavior classifier performance for lunge (A) and unilateral wing extension (B). Three twenty-minute representative recordings of male-male pairs for each species were manually scored for both actions (expert annotation, same pairs as in Fig. 1A) and exposed to automated behavior classifiers trained previously for *D. melanogaster* (*57*), implemented in JAABA (*55*). Manual (x-axis) vs. classifier-generated annotations (y-axis) are compared in log_10_ scale, using either the number of bouts (lunge) or their cumulative time duration (UWE). Solid lines, diagonals; dashed lines, linear fits. Adjusted R^2^, significance by F-tests for fits, and Pearson correlations (r) indicated at bottom right. *D. melanogaster*, black; *D. santomea,* orange; *D. yakuba*, green; *D. simulans*, *D. mauritiana*, *D. sechellia*, *D. erecta*, gray. Note near-perfect overlap with the diagonal for lunge and strong correlation for UWE despite classifier false positives in cases with few manual annotations (all subgroup species except *D. santomea* show very little male-male courtship). **(C,D)** Aggression (lunge, C) and courtship (UWE, D) measurements from automated classifiers in all recorded male-male pairs for *melanogaster* subgroup species and additional *D. santomea* strains. *D. santomea* 00 (also called STO.4) is the strain included in the behavioral screen and used throughout as “wildtype” (National Drosophila Species Stock Center 14021.0271.00). Strains 146 (STO-CAGO 1482), 148 (STO.7), and 151 (STO.6) also derive from São Tomé from females collected between 1100 and 1500 m elevation (*117*). Boxplots show full distribution range within whiskers (excluding statistically identified outliers) and second and third quartiles within boxes, with medians in bold. Individual data points (including outliers) overlayed as gray dots. Outliers retained in all summary metrics and statistical comparisons here and throughout. Lettered statistical groupings assigned by post-hoc Dunn’s tests following significant Kruskal-Wallis. Note low aggression and high courtship conserved in *D. santomea* strains (orange), low aggression and courtship in the sibling species *D. yakuba* (green), and high aggression and low courtship in *D. melanogaster* (black). **(E,F)** Summaries of penetrance (x-axes) and expressivity (y-axes) for aggression (E) and courtship (F) across subgroup species and *D. santomea* strains. Metrics derived from data in (C,D).

**Figure S3.**
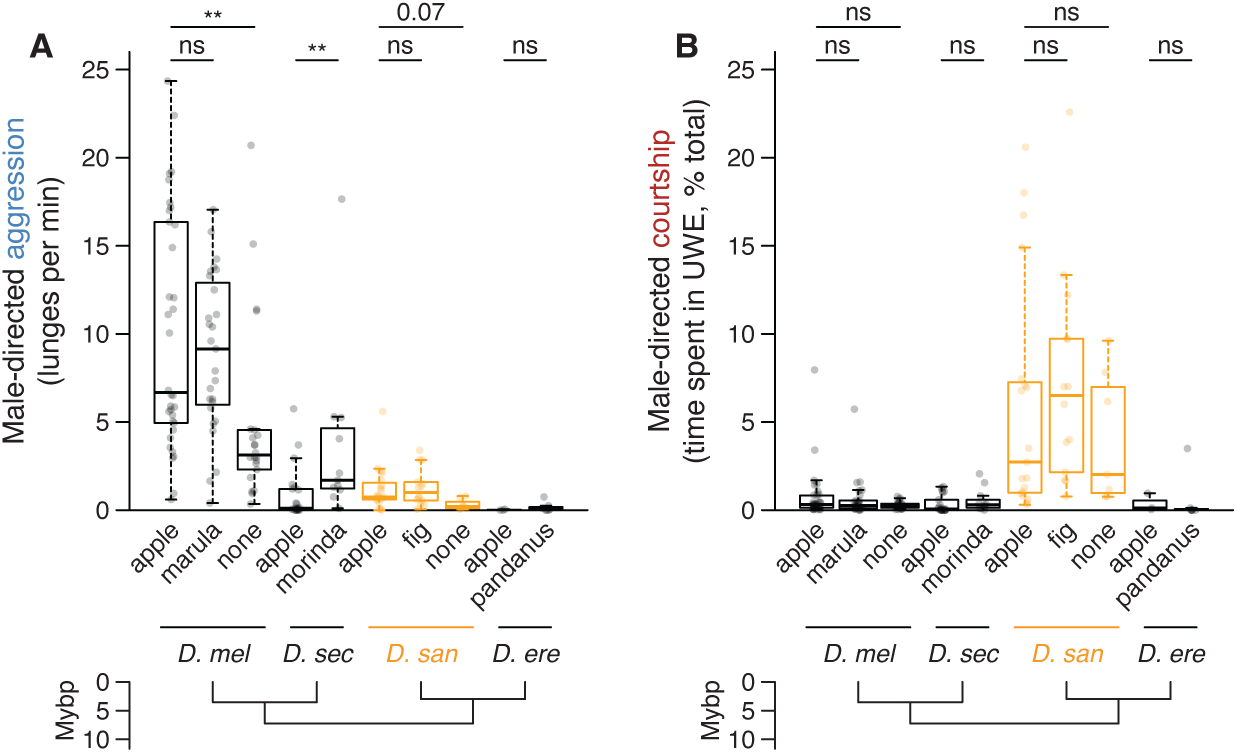
Intermale aggression and courtship in *melanogaster* subgroup species with fruit host specializations. **(A,B)** Aggression (A) and courtship (B) in *D. melanogaster*, *D. sechellia*, *D. santomea* (orange), and *D. erecta* male-male pairs competing over the standard apple juice substrate and other fruits based on reported specializations (*54*). Significance for each fruit/species combination by pairwise Mann-Whitney *U* tests to apple. Removing the food source significantly reduces aggression in *D. melanogaster* (*51*). *Morinda* (also called noni) significantly increases aggression but not courtship among *D. sechellia* males. *D. santomea* aggression and courtship depend only weakly if at all on food presence and source. *Morinda* and fig juices obtained commercially, *Marula* (*Sclerocarya birrea*) and *Pandanus* (*Pandanus furcatus*) obtained from botanical gardens (table S1).

**Figure S4.**
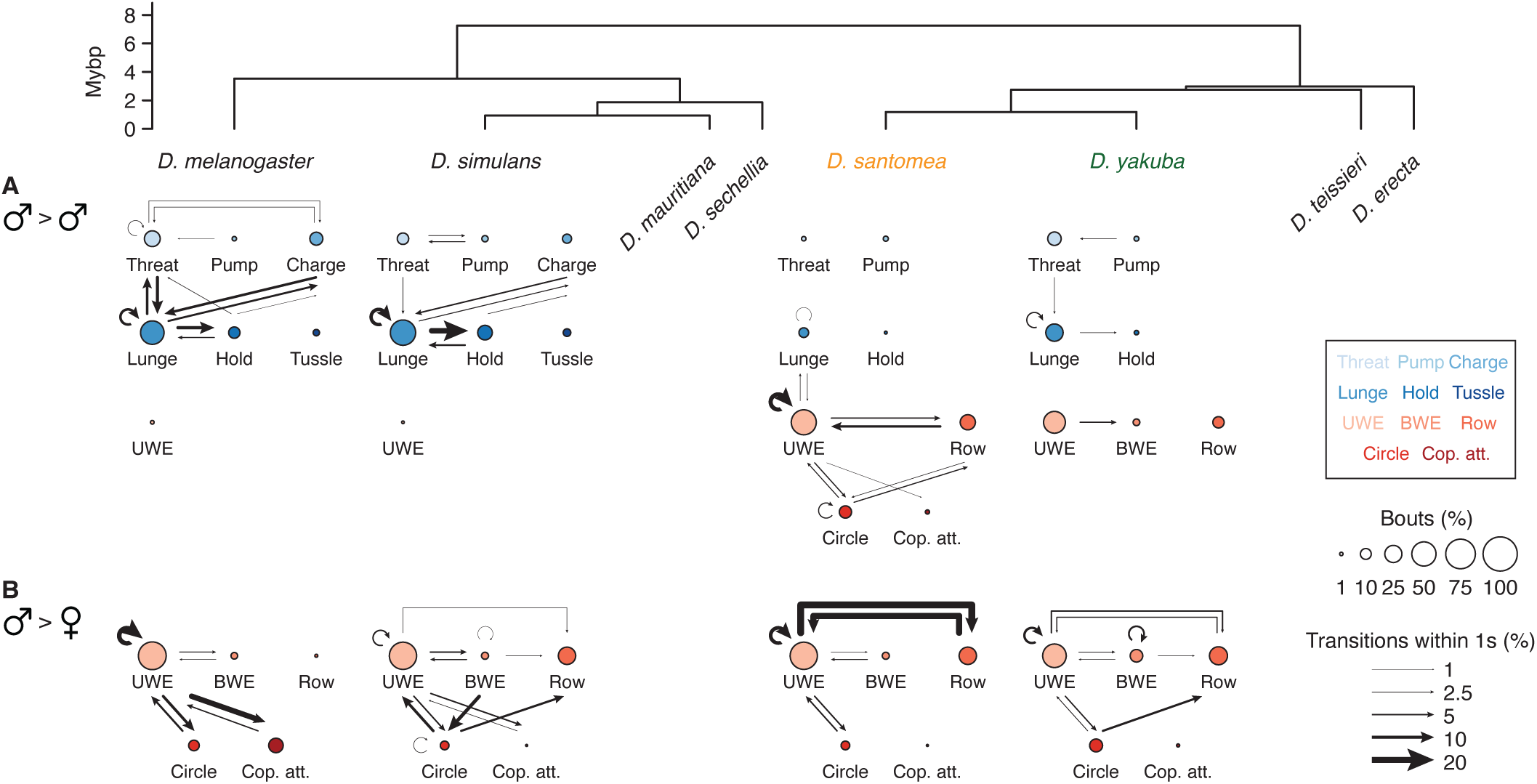
Male- and female-directed social behaviors by *D. melanogaster*, *D. simulans*, *D. santomea,* and *D. yakuba* males. **(A,B)** Ethograms representing male-male (A) and male-female (B) social behaviors for *melanogaster* subgroup species of interest. Phylogeny also includes *D. mauritiana* and *D. sechellia* from the *simulans* clade, *D. teissieri* from the *yakuba* clade, and *D. erecta* without associated ethograms. Female-directed ethograms represent expert annotation for three recordings of male-female pairs per species, with male flies similarly prepared by single-housing and interactions taking place under identical conditions. Male-directed ethograms reproduced from Fig. 1A for visual comparison to female-directed counterparts. Note nearly exclusive male-directed aggressive actions (blue nodes) by *D. melanogaster* and *D. simulans* males but prevalence of male-directed courtship actions (red nodes) by *D. santomea* and *D. yakuba* (though in *D. yakuba* total male-direcetd actions are many fewer on average than *D. santomea)*, matching the actions taken when paired with females.

**Figure S5.**
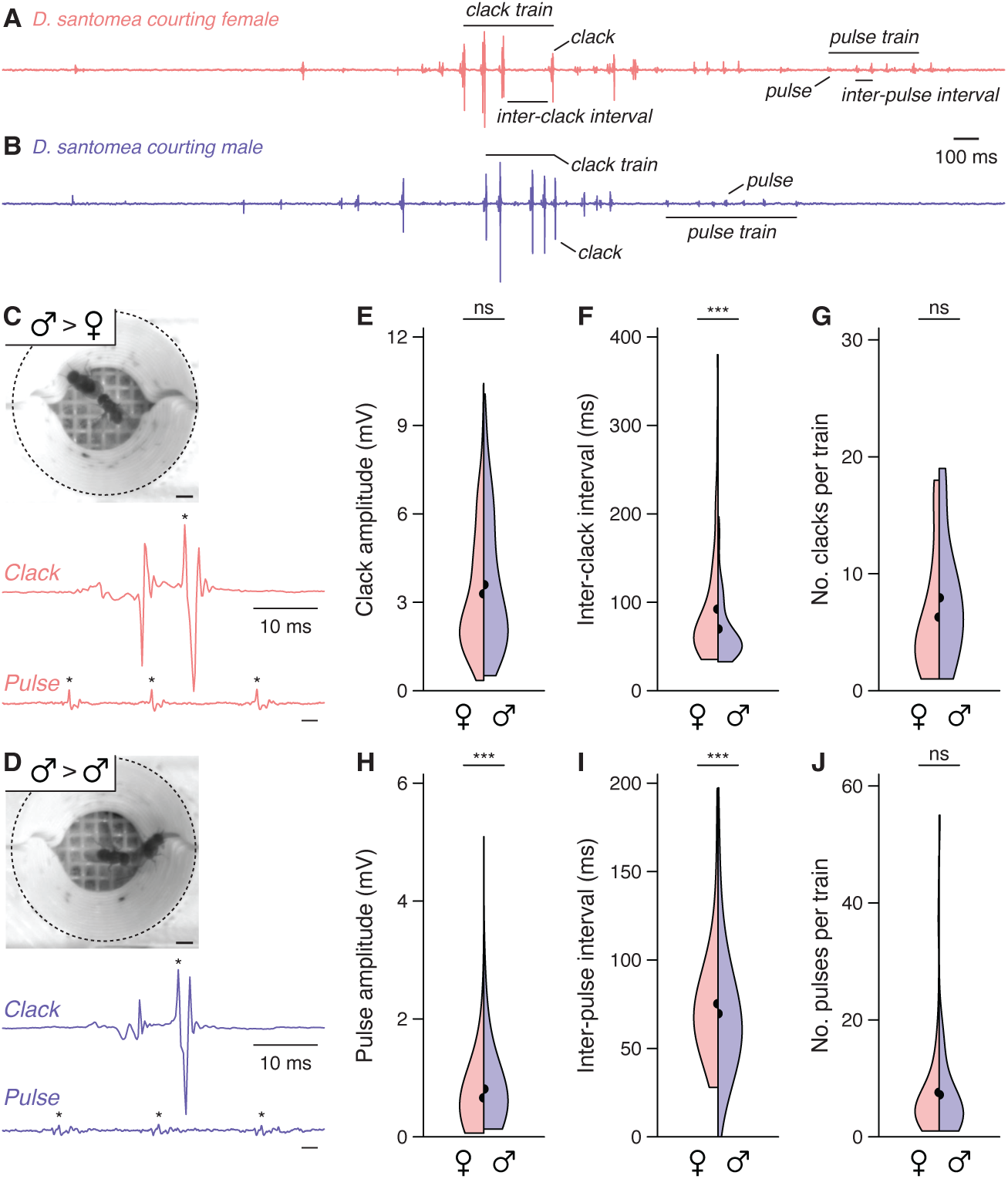
Acoustic features of female- and male-directed courtship song by *D. santomea* males. **(A,B)** Representative courtship song traces by *D. santomea* males toward a female (A, pink) or male conspecific target fly (B, purple). Key acoustic features are indicated including pulse, pulse train, inter-pulse interval (IPI), clack, clack train, and inter-clack interval (ICI) (*63*). Traces reproduced and enlarged from Fig. 1I,J. **(C,D)** Video stills of song recording chambers (*60*) containing *D. santomea* males courting either a female (C) or male (D). Isolated waveforms of single clacks and pulse trios shown below, with asterisks indicating event peaks. Scale bars, 1 mm. **(E-J)** Quantitative acoustic analyses of female-(pink) and male-directed *D. santomea* courtship songs (purple). Clacks (E-G) and pulses (H-J) were identified by expert annotation from three recordings with each sex (clack, 32-103 events per recording; pulse, 140-346 events per recording). Data shown as “hemiviolin” plots in which female- and male-directed distributions are vertically mirrored. Distribution means are semicircles on the appropriate side. Note general similarity between female- and male-directed song features (significant differences based on Mann-Whitney *U* tests despite small difference magnitudes are often because so many data points make up the underlying distributions). Analyses of absolute event abundances were omitted since annotations captured only a fraction of events in each recording.

**Figure S6.**
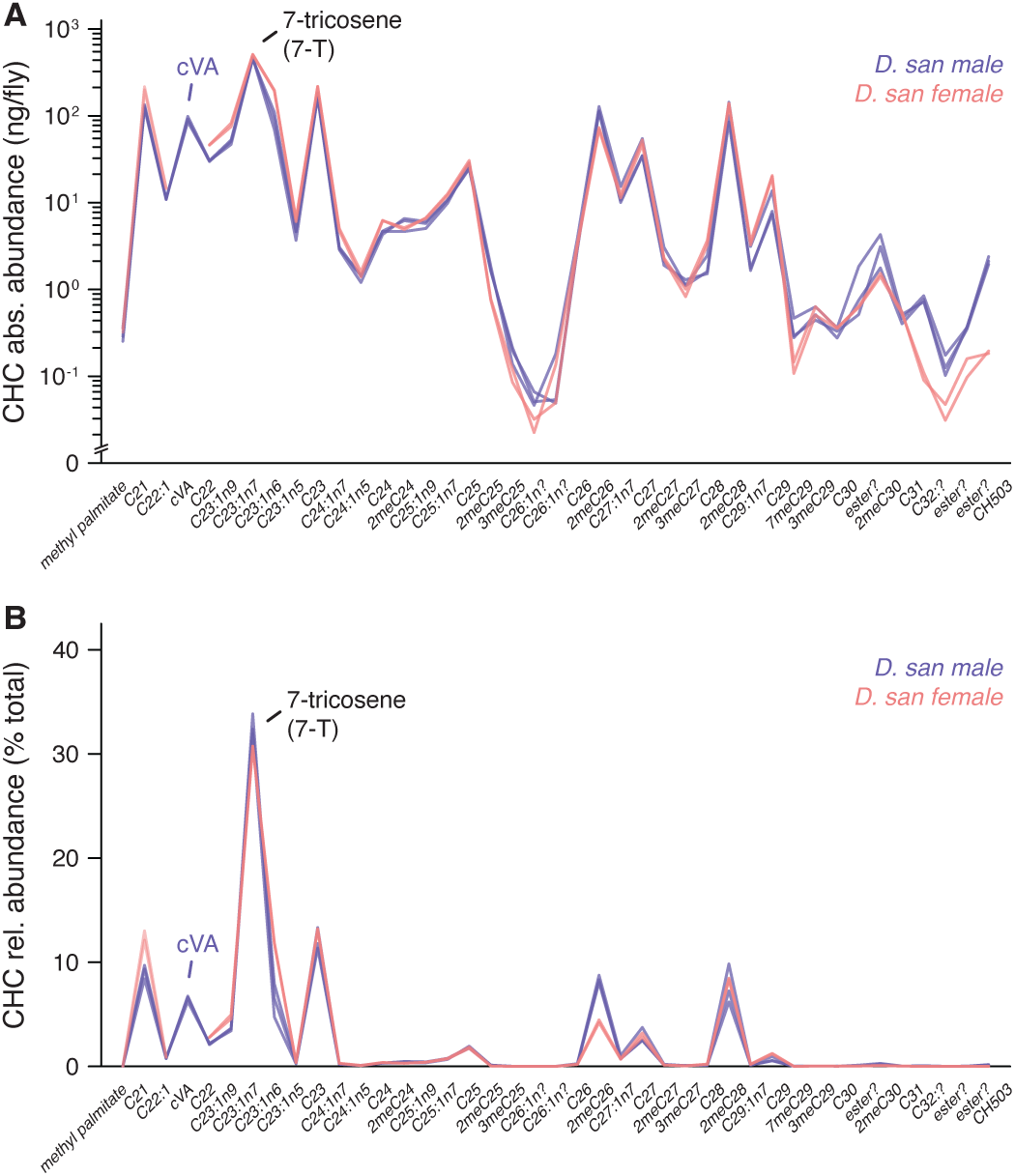
GC-MS profiling of *D. santomea* male and female cuticular hydrocarbons (CHCs) **(A,B)** Absolute (A) and relative abundances (B) of 41 compounds identifiable as peaks in *D. santomea* male (purple) and female hexane extracts (pink). Individual measurements are made on ten pooled, age-matched adult flies (3 male, 2 female replicates). Calculation of absolute abundance uses known concentration of the internal standard (octadecane/C18, not shown) and normalizes for number of flies included in the extract. Relative abundances normalize to the summed absolute abundance of all compounds excluding the standard. Male-specific (Z)-11-octadecenyl acetate (cVA, purple) and monomorphic (Z)-7-tricosene (7-T, black) are indicated. Question marks in compound names indicate uncertainty in chemical assignment. See table S5 for retention times and diagnostic ions used to identify and assign CHCs.

**Figure S7.**
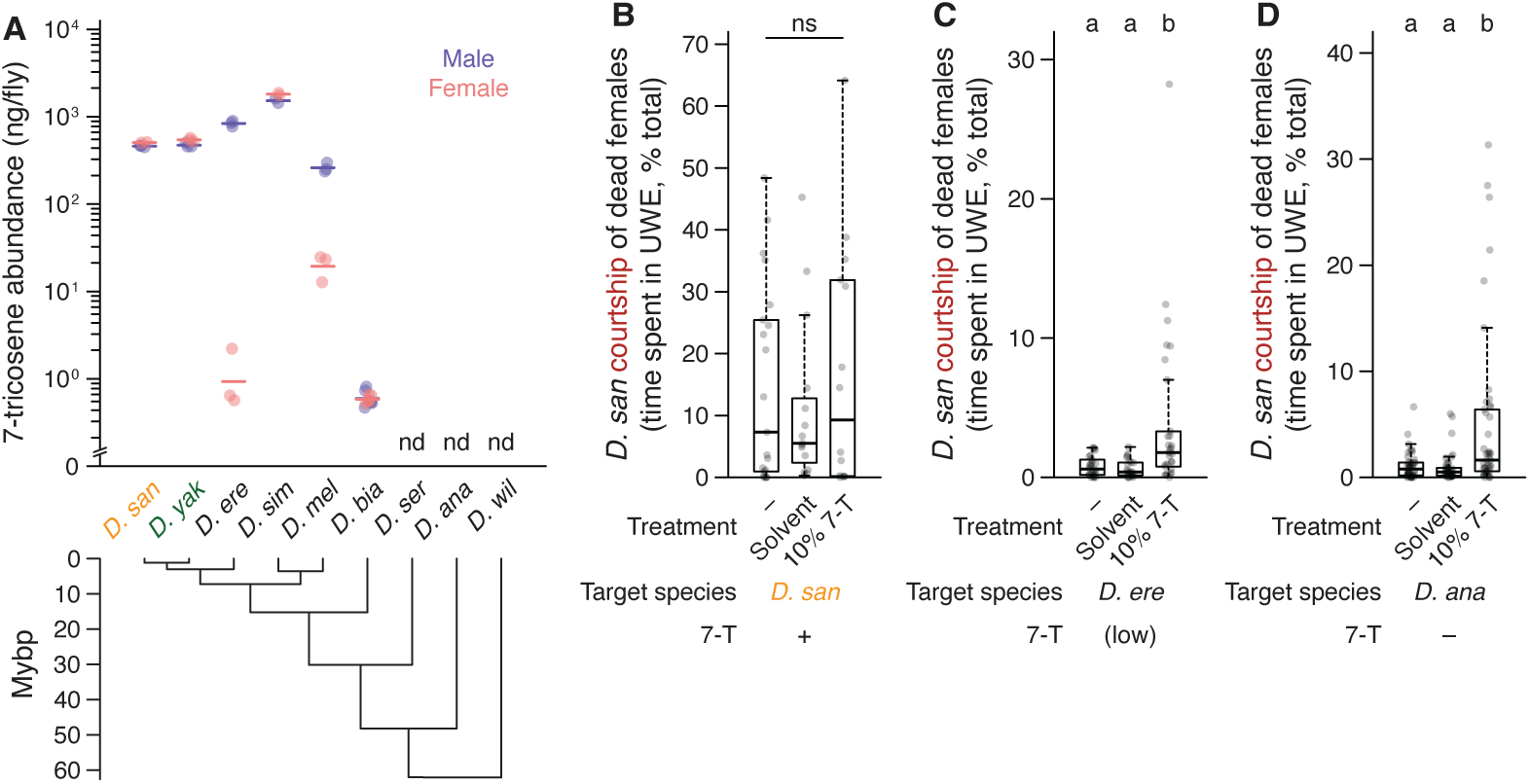
7-tricosene (7-T) abundance in males and females of various *Drosophila* species and additional 7-T add-back experiments. **(A)** Absolute 7-T abundances on males (purple) and females (pink) *Drosophila* measured by GC-MS on hexane extracts from ten pooled, age-matched adult flies (2-8 replicates each). Monomorphic species (*D. santomea, D. yakuba*, *D. simulans*) show similar 7-T abundance between sexes while dimorphic species show large differences (*D. erecta*, *D. melanogaster*). nd, not detected in either sex (*D. serrata*, *D. ananassae*, *D. willistoni*). **(B-D)** Fraction of time *D. santomea* males spend courting conspecific (B) or heterospecific dead females (C,D) perfumed with 10% 7-T (20 µg/fly) or solvent control (hexane). Heterospecific female targets selected for having either low (*D. erecta*) or undetectable (*D. ana*) endogenous 7-T levels (from GC-MS, A). *D. santomea* female targets as positive controls. 7-T addition elicits courtship from *D. santomea* males in both heterospecific cases. Significance by Dunn’s tests.

**Figure S8.**
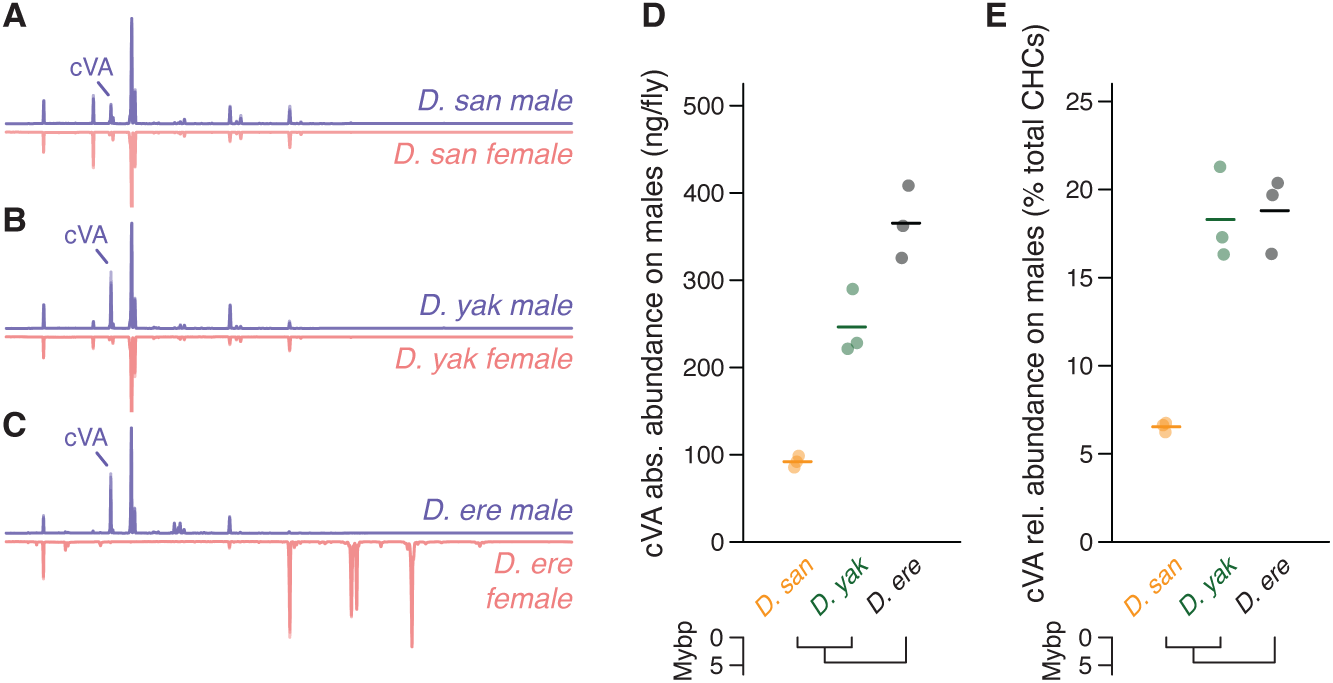
Reduced cVA on *D. santomea* males compared to *D. yakuba* and *D. erecta*. **(A-C)** Mirrored gas chromatograms representing male (purple, top) and female (pink, bottom) cuticular hydrocarbon (CHC) profiles of *D. santomea* (A), *D. yakuba* (B), and *D. erecta* (C). Individual measurements made on hexane extracts from ten pooled, age-matched adult flies (2-3 replicates each). Peaks representing male-specific (Z)-11-octadecenyl acetate (cVA) indicated. *D. santomea* chromatograms reproduced from Fig. 2A for visual comparison. **(D,E)** Absolute (D) and relative abundances (E) of cVA on *D. santomea* (orange), *D. yakuba* (green), and *D. erecta* males. Calculation of absolute abundance uses known concentration of the internal standard (C18) and normalizes for number of flies included in the extract. Relative abundances normalize to the summed absolute abundance of all compounds detected excluding the standard. *D. santomea* males show 63% and 75% reduced cVA comparing absolute abundance means or 64% and 65% reductions comparing relative abundance means to *D. yakuba* and *D. erecta*, respectively.

**Figure S9.**
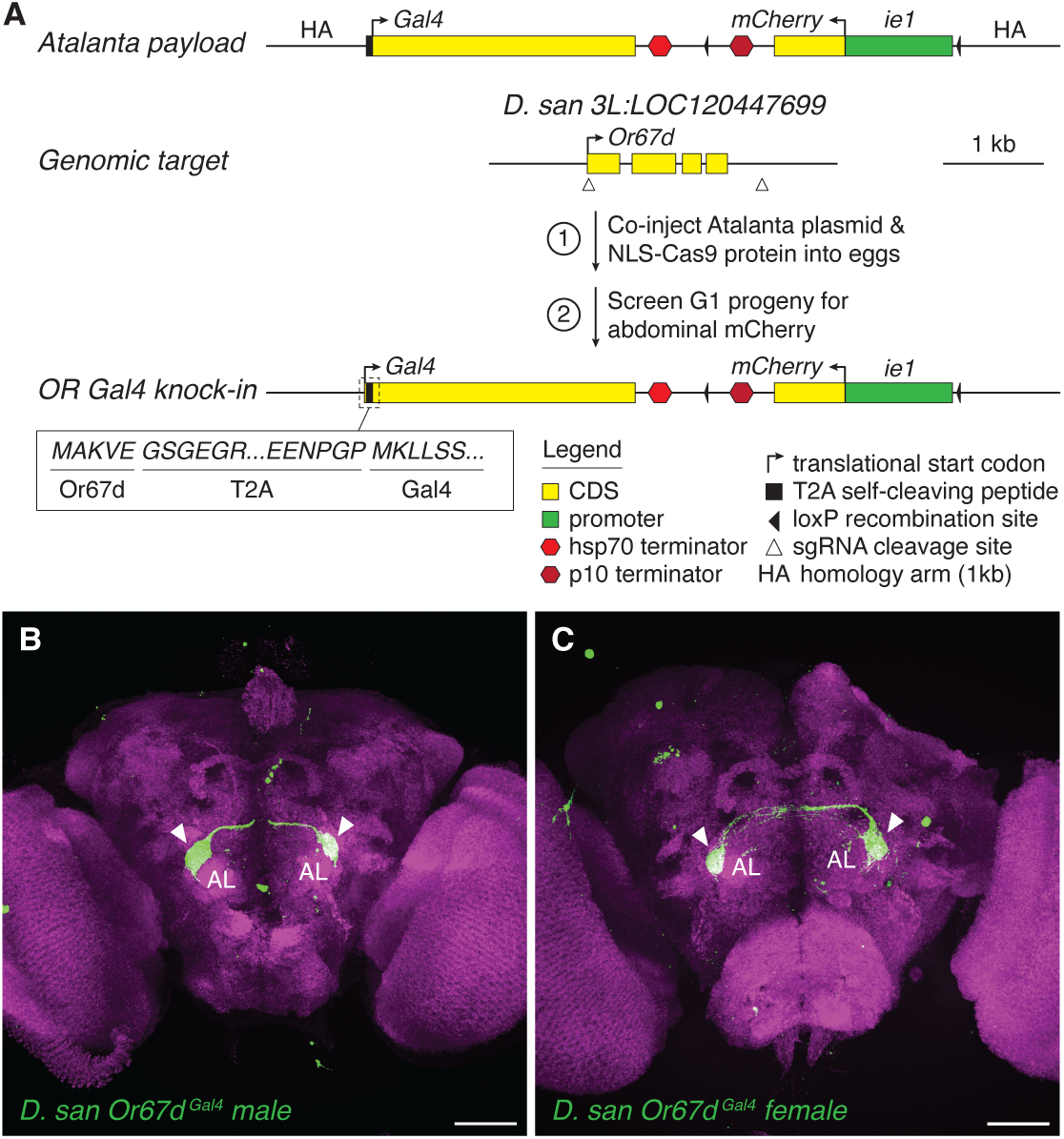
CRISPR-Cas9 HDR knock-in strategy used to generate *D. santomea Or67d^Gal4^*. **(A)** Schematic diagrams of the Gal4 transgene and selection marker assembled into an Atalanta vector (pJAT32, addgene #204297) (*158*) and *Or67d* genomic target locus shown with upstream and downstream sgRNA cleavage sites. Transgene (above) contains two cassettes: *T2A-Gal4-stop(hsp70)* oriented natively to the genomic target, and downstream *ie1-mCherry-stop(p10)* fluorescent marker for successful integration (*32*) with inverted orientation. One kilobase homology arms flank both ends. *Or67d* genomic locus (below) is targeted using sgRNAs directing double-strand breaks to immediately (15 bp) downstream of the translational start codon and 348 bp downstream of the stop, resulting in removal of 1152 of 1167 protein coding base pairs (99%). Co-injection of the assembled Atalanta vector and nls-Cas9-nls protein into wildtype *D. santomea* eggs gives mosaic *ie1-mCherry^pos^* G0 progeny carrying *T2A-Gal4* inserted in-frame with the five residual residues of *Or67d*. G1 progeny resulting from G0 crosses to wildtype are re-screened for abdominal *ie1-mCherry* expression to ensure germline transmission. In this strategy spatiotemporal patterns and magnitude of Gal4 expression are controlled entirely by *Or67d* endogenous genomic regulation. **(B,C)** Photomicrographs showing labeling patterns of *Or67d^Gal4^* in the whole brain of a *D. santomea* male (B, same fly from which close-up of antennal lobes (ALs) is shown in Fig. 3F) and female (C). Immunostaining for Gal4-dependent cytoplasmic tdTomato pseudocolored green with Bruchpilot synaptic counterstain (using nc82 monoclonal antibody) in magenta. Uniglomerular labeling pattern in each AL (arrowheads) matches the expected size and position of cVA-sensitive DA1 targeted by Or67d OSNs (*145*). Bifurcating OSN commissures (axon bundles) from the antennae leading into ALs can also be seen. Scale bars, 50 µm.

**Figure S10.**
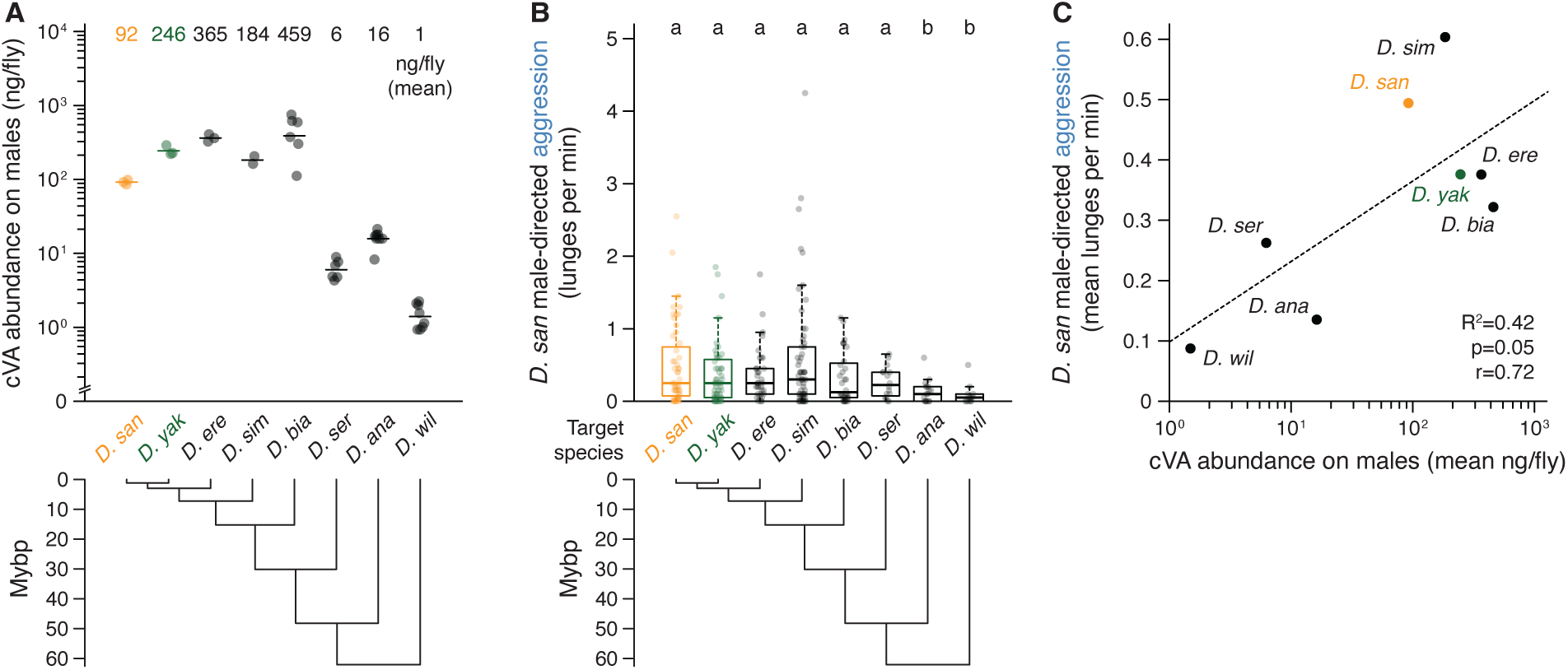
Ineffectiveness of naturally varying cVA levels on various species to promote aggression by *D. santomea* males. **(A)** Absolute cVA abundance on *Drosophila* males measured by GC-MS on hexane extracts from ten pooled, age-matched adult flies (2-8 replicates each). Calculation uses known concentration of the internal standard (C18) and normalizes for number of flies included in the extract. *D. santomea* (orange), **(A)** *D. yakuba* (green), and *D. erecta* measurements reproduced from fig. S8 for visual comparison to additional species. Means indicated above and phylogeny below. **(B)** Spontaneous aggression (lunges per minute) exhibited by *D. santomea* males toward conspecific (orange) or heterospecific males (*D. yakuba* green, all others black) in single-housed pairs during twenty-minute interactions. Statistical groupings by Dunn’s tests shown above and phylogeny below. Note similarly low or even decreased attack levels toward males of all species. **(C)** Correlation between cVA abundance measured by GC-MS and aggression elicited from *D. santomea* males. Dashed line, linear fit. Adjusted R^2^, significance by F-test for fit, and Pearson correlation (r) indicated at bottom right. Note marginally significant correlation driven mostly by low attack rates toward distant species (*D. serrata*, *D. ananassae*, *D. willistoni*) with many other pheromone changes in addition to low cVA. Three of four species with higher cVA abundance than *D. santomea* elicit less attack than observed among *D. santomea* males (*D. simulans* is the exception but the slight increase is non-significant). Data derived from (A,B).

**Figure S11.**
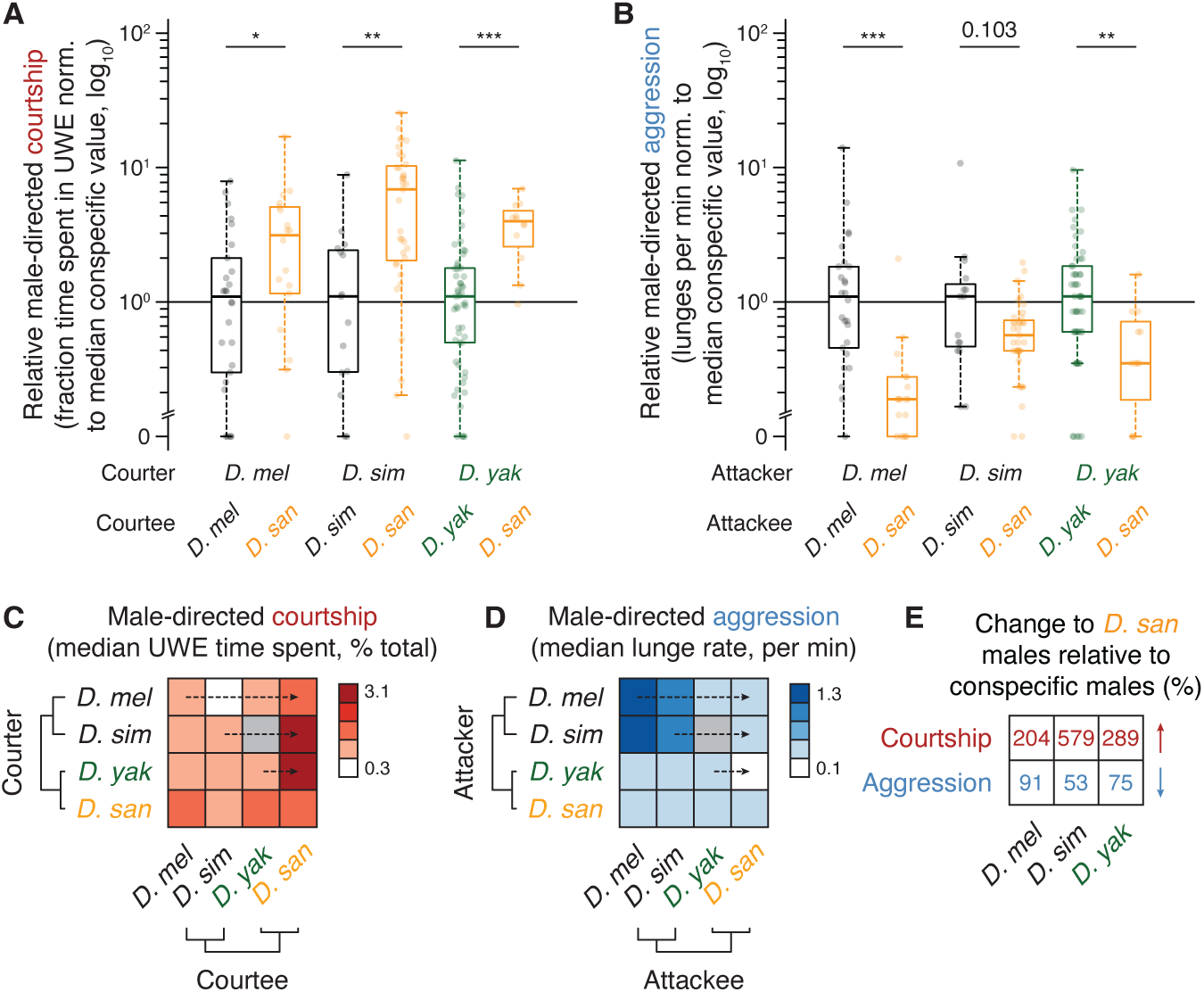
Social behavior changes by *melanogaster* subgroup males toward *D. santomea* males. **(A)** Relative fraction of time *D. melanogaster*, *D. simulans*, and *D. yakuba* (green) males court conspecific males (first distribution of each pair) vs. *D. santomea* males (second distribution, orange). All measurements within each species normalized to the conspecific median and log_10_ transformed. Males of all three *melanogaster* subgroup species show significantly increased courtship toward *D. santomea* males by Mann-Whitney *U* tests, including *D. melanogaster* where 7-T (abundant on *D. santomea* males) is male-specific and sexually aversive (*71–74*). **(B)** Similar to (A) but showing paired relative distributions for aggression within and between males of the three subgroup species. Note decreases in male-directed aggression by all three toward *D. santomea* (significant for *D. melanogaster* and *D. yakuba* by Mann-Whitney *U* tests). **(C)** Heatmap showing median fraction time spent courting by males in conspecific (diagonal) and heterospecific male-male pairs (off-diagonal) among the four subgroup species. Rows (“courters”) are males whose courtship is measured (UWE). Columns (“courtees”) are males targeted by courtship. Gray entry where data not collected. Data derived from (A) and additional pairings. Horizontal dashed arrows identify relevant comparisons between conspecific male- and *D. santomea* male-directed courtship for visual aid. **(D)** Similar heatmap showing median aggression (lunge rate) by male “attackers” (rows) and “attackees” (columns). Data derived from (B) and additional pairings. Note consistently decreased aggression toward *D. santomea* males. **(E)** Summarized changes in courtship (first row) and aggression rates (second row) by heterospecific males toward *D. santomea* compared to conspecific rates for each, derived from data in (A,B). Percent increases in first row (courtship) and percent decreases in second (aggression).

**Figure S12.**
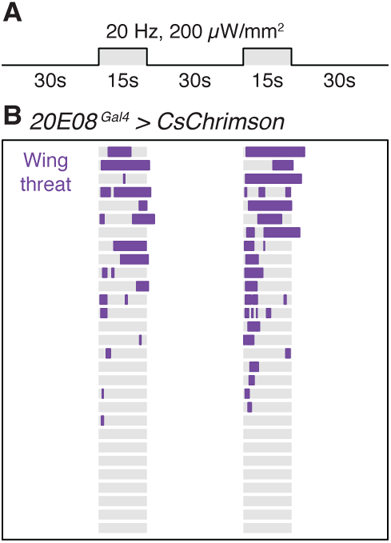
Induction of time-locked wing threat by photoactivation of AIP neurons in solitary *D. santomea* males. **(A)** Optogenetic photoactivation scheme for solitary *D. santomea* males carrying *20E08^Gal4^* and Gal4-dependent CsChrimson. Group-housed *20E08^Gal4^* tester males or genetic controls are exposed to two fifteen-second photostimulation (PS) blocks with a thirty-second baseline and inter-block interval. **(B)** Manually scored wing threat rasters (purple) for solitary *20E08^Gal4^* males during the two-minute trial. Note threat induction time-locked to PS in most flies. Genetic controls showed no threat.

**Figure S13.**
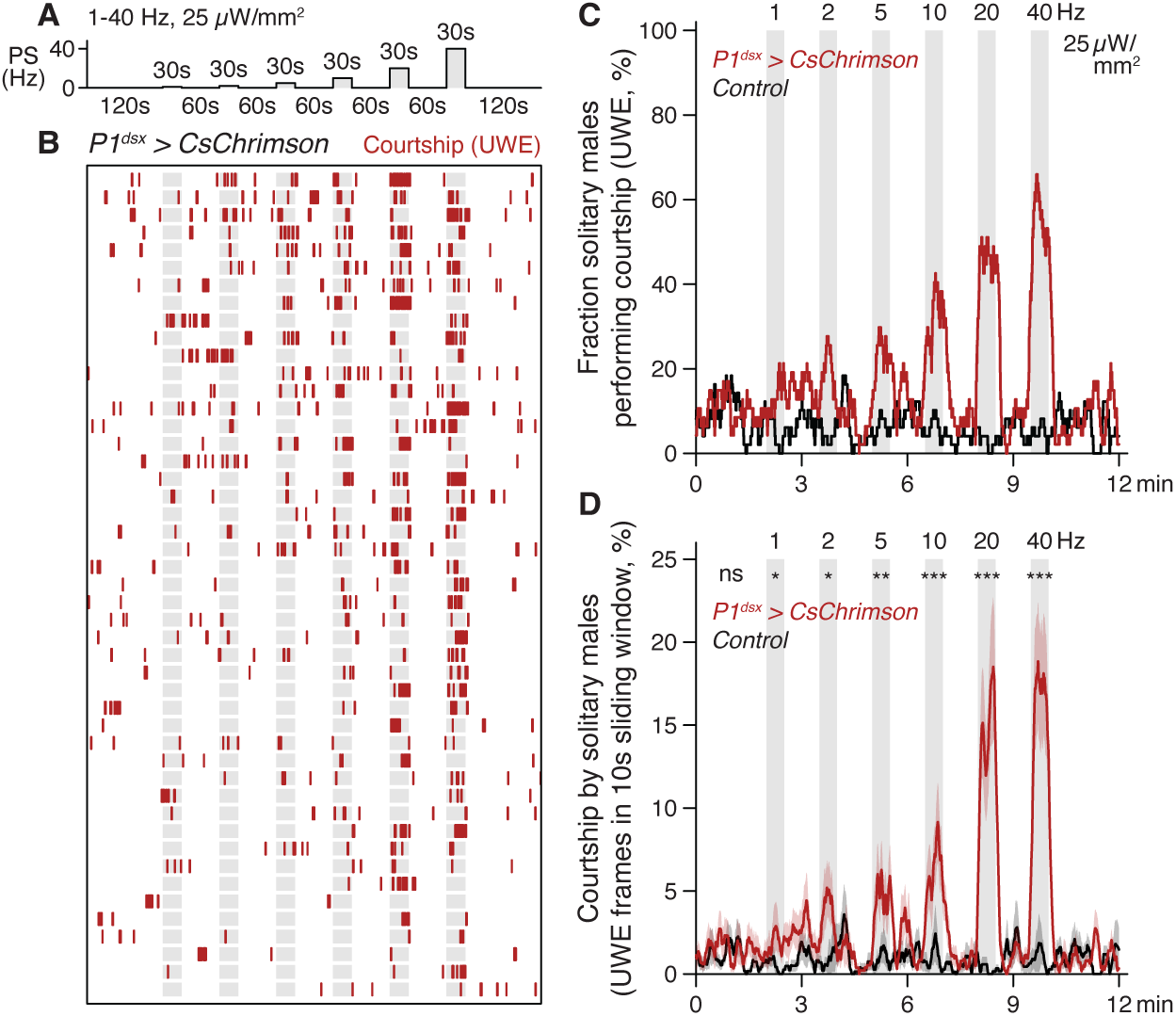
Induction of time-locked courtship by photoactivation of P1^dsx^ neurons in solitary *D. santomea* males. **(A)** Optogenetic photoactivation scheme for solitary group-housed *D. santomea* males carrying *P1^dsx^* split-Gal4 (*71G01^DBD^;dsx^AD^*) and Gal4-dependent CsChrimson. *P1^dsx^* tester males or a genetic control are exposed to six thirty-second photostimulation (PS) blocks with fixed intensity and monotonically increasing frequency as indicated, with a two-minute baseline and one-minute inter-block intervals. **(B)** Courtship rasters (red) for *P1^dsx^* tester males during the twelve-minute trial. Note early but sporadic courtship bouts that time-lock to PS blocks and increase in penetrance starting at 5 and 10 Hz. **(C,D)** Penetrance (C) and expressivity (D) of courtship (UWE) elicited by *P1^dsx^* PS. Significant courtship expression during PS blocks compared to the genetic control in (D) by Mann-Whitney *U* tests.

**Figure S14.**
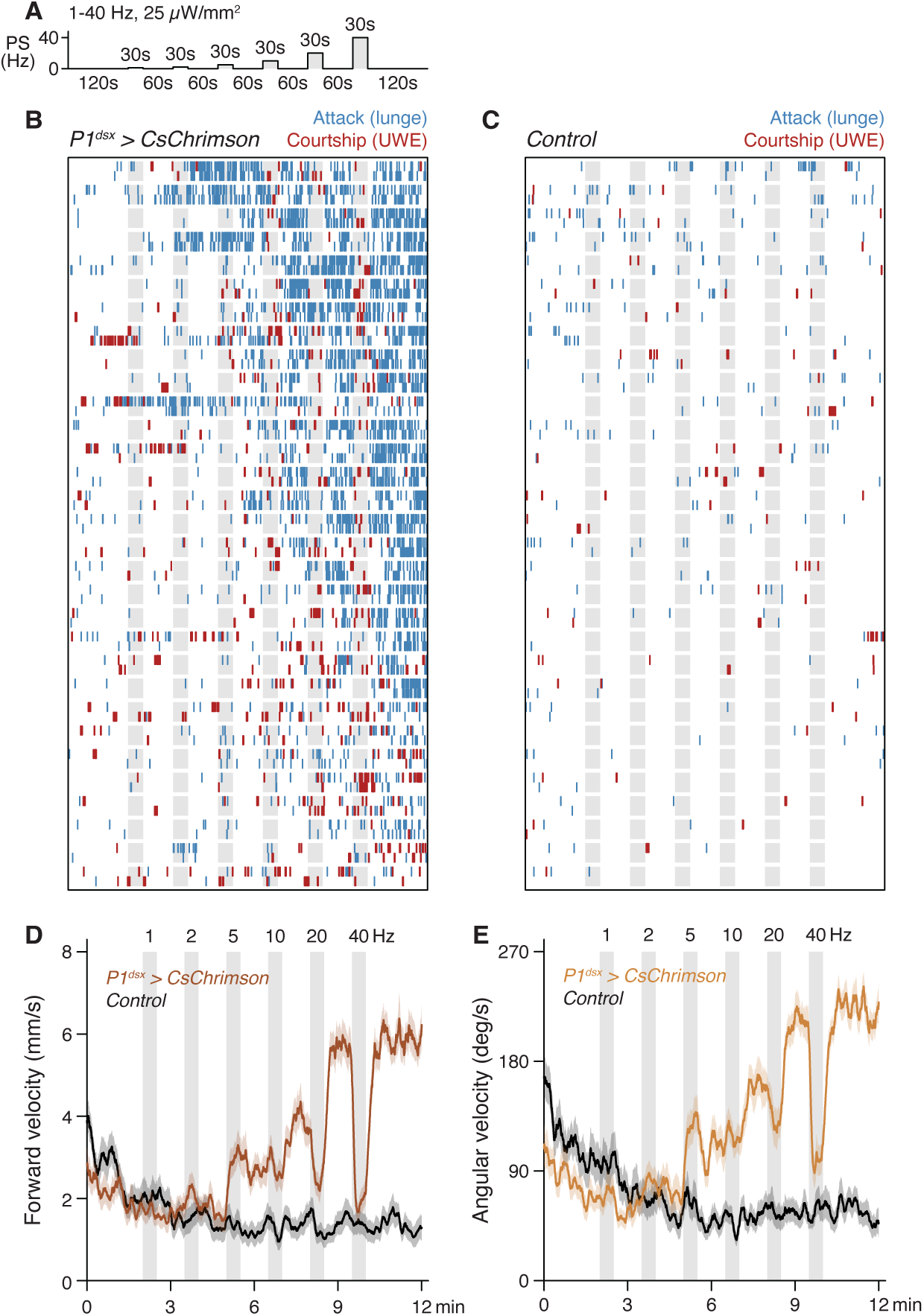
Induction of courtship and attack by photoactivation of P1^dsx^ neurons in same-genotype pairs of *D. santomea* males. **(A)** Optogenetic photoactivation scheme for same-genotype pairs of group-housed *D. santomea* males carrying *P1^dsx^* split-Gal4 and Gal4-dependent CsChrimson. *P1^dsx^* tester males or a genetic control are exposed to six thirty-second photostimulation (PS) blocks with fixed intensity and monotonically increasing frequency as indicated, with a two-minute baseline and one-minute inter-block intervals. **(B,C)** Behavior rasters for *P1^dsx^* male pairs (B) and genetic controls (C) during the twelve-minute trial. Blue, attack (lunge); red, courtship (UWE). Note early induction of attack which increases in penetrance and expressivity as the trial continues. Attacks interrupted by time-locked courtship during later PS blocks (>5 Hz). Genetic controls show little courtship or attack. **(D,E)** Locomotor dynamics in *P1^dsx^* males (browns) and genetic controls (black). Per-frame calculations of each fly’s forward velocity (D) and angular velocity (E) derived from automated tracking (*56*). Data smoothed by averaging within a ten-second sliding window. Envelopes represent s.e.m. PS time-locked locomotor arrest similar to observed previously in *D. melanogaster* (*57*).

**Figure S15.**
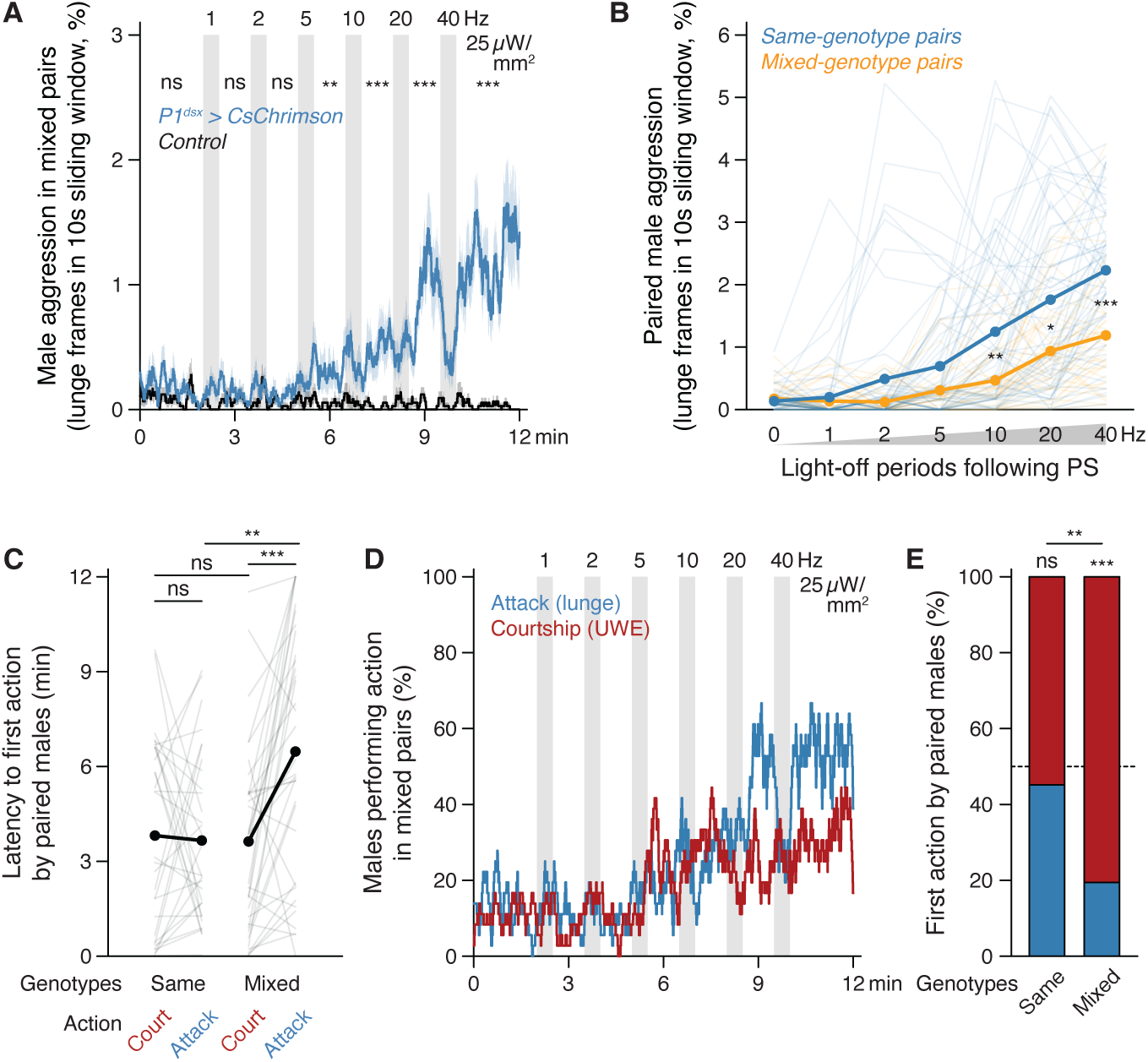
Photoactivation effects of *D. santomea* P1^dsx^ neurons in mixed-genotype male pairs. **(A)** Expressivity and temporal dynamics of attack (lunge) elicited by PS in mixed-genotype pairs between a *P1^dsx^* split-Gal4 tester male carrying Gal4-dependent CsChrimson (or genetic control) and group-housed wildtype *D. santomea* target male. Traces show the mean fraction of frames containing lunge in a ten-second sliding window with s.e.m. envelopes for *P1^dsx^*(blue) and control (black). PS block frequency, intensity, and timing indicated above for the twelve-minute trial. Note significant attack induction (Mann-Whitney *U* tests) in *P1^dsx^* starting after 5 Hz. **(B)** Attack elicited during inter-bout intervals and after final PS in same-genotype *P1^dsx^* male pairs (blue, data from Fig. 4I) vs. mixed-genotype pairs between a *P1^dsx^*tester and group-housed wildtype male (orange, from A). Period means connected with solid lines and data from individual flies shown as thin lines. Note significant right shift in mixed pairs (Mann-Whitney *U* tests) indicating higher PS frequencies required to induce intense attack. **(C)** Action latencies for courtship and attack exhibited by *P1^dsx^* tester males in same- and mixed-genotype pairs under identical PS trial structures. Data points derived from the same fly pair (or tester fly for mixed pairs) are connected as lines and distribution means shown as paired dots. Attack and courtship latencies are similar on average in same-genotype pairs whereas attack is significantly delayed in mixed pairs. Significance by paired or unpaired Mann-Whitney *U* tests (for within- or between-genotype group comparisons, respectively). **(D)** Fraction of flies showing attack (lunge, blue) and courtship (UWE, red) in *P1^dsx^* mixed-genotype pairs. **(E)** Relative frequencies of *P1^dsx^* same- and mixed-genotype pairs showing courtship (red) or attack (blue) first. Significance between distributions and between each distribution and a null probability of 50% (random chance) determined by binomial tests. Note courtship precedence in mixed pairs.

**Figure S16.**
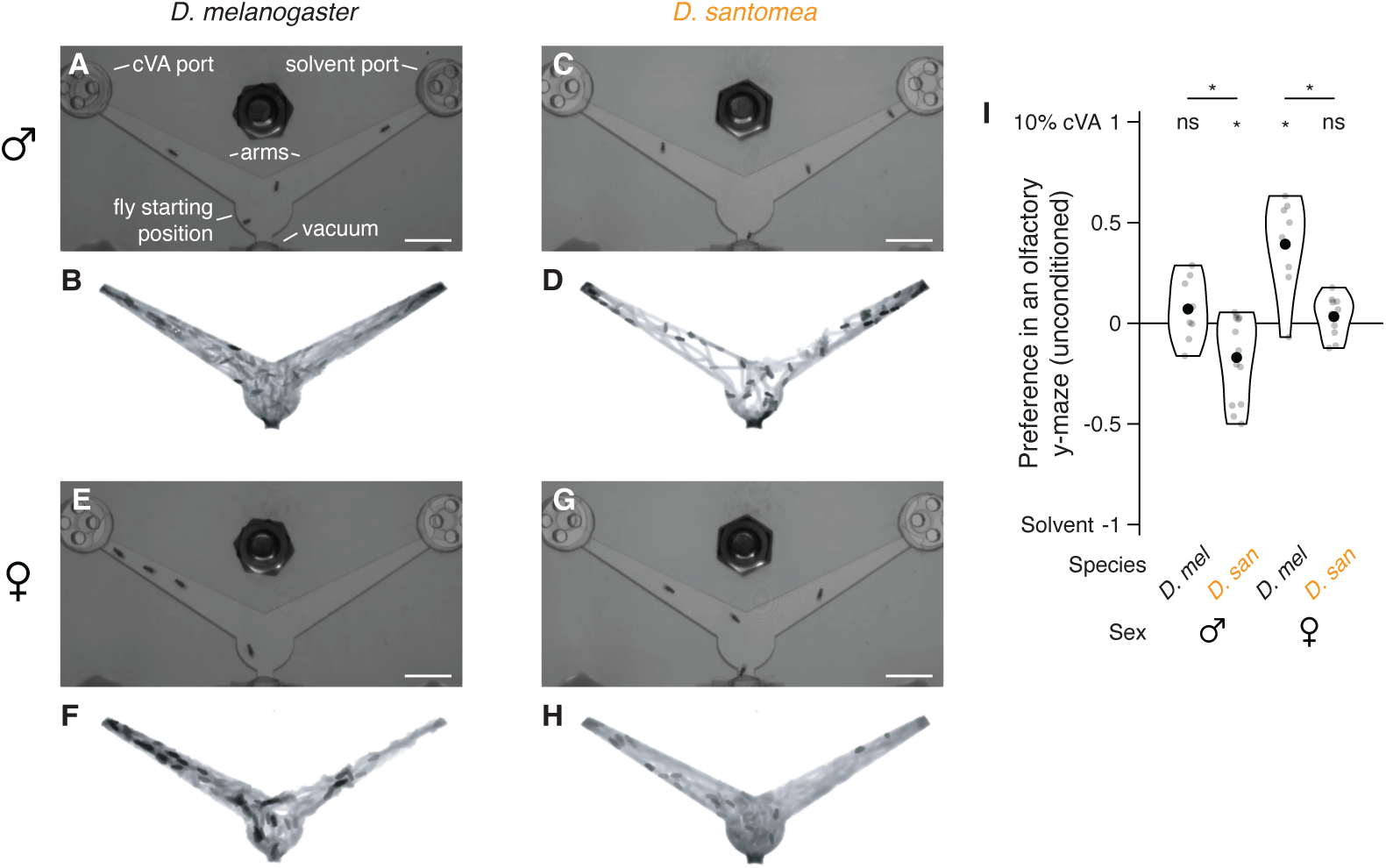
Unconditioned place preference tests with cVA in *D. melanogaster* and *D. santomea*. **(A)** Video still illustrating the y-maze arena. Four group-housed *D. melanogaster* males are introduced into the central bowl at the bottom and odors (10% cVA in acetone or acetone control) into ports at the head of each arm. Vacuum pressure continuously pulls air down the arms through an outlet. Flies can walk freely within the bowl and arms during ten-minute trials and odors are replaced into the same port (to avoid contamination) before each trial. Scale bar, 10 mm. **(B)** Cumulative position trace representing movement of all four flies throughout the duration of the trial. Position density in each of multiple trials is used to calculate the odor preference index. **(C-H)** Video stills and cumulative position traces as in (A,B) for *D. santomea* males (C,D), *D. melanogaster* females (E,F), and *D. santomea* females (G,H). Scale bars, 10 mm. **(I)** Odor preference indices by species and sex. Index calculated as the difference in fly occupancy between the two arms normalized by the sum. Small gray dots represent single trials with 2-4 flies each (8-14 trials and 26-44 flies per species/sex). Distributions represented as violins with means shown as large black dots. Note significantly reduced attraction (or increased aversion) to 10% cVA in both *D. santomea* males and females (orange) compared to *D. melanogaster* (black). Significance by Mann-Whitney *U* tests.

**Figure S17.**
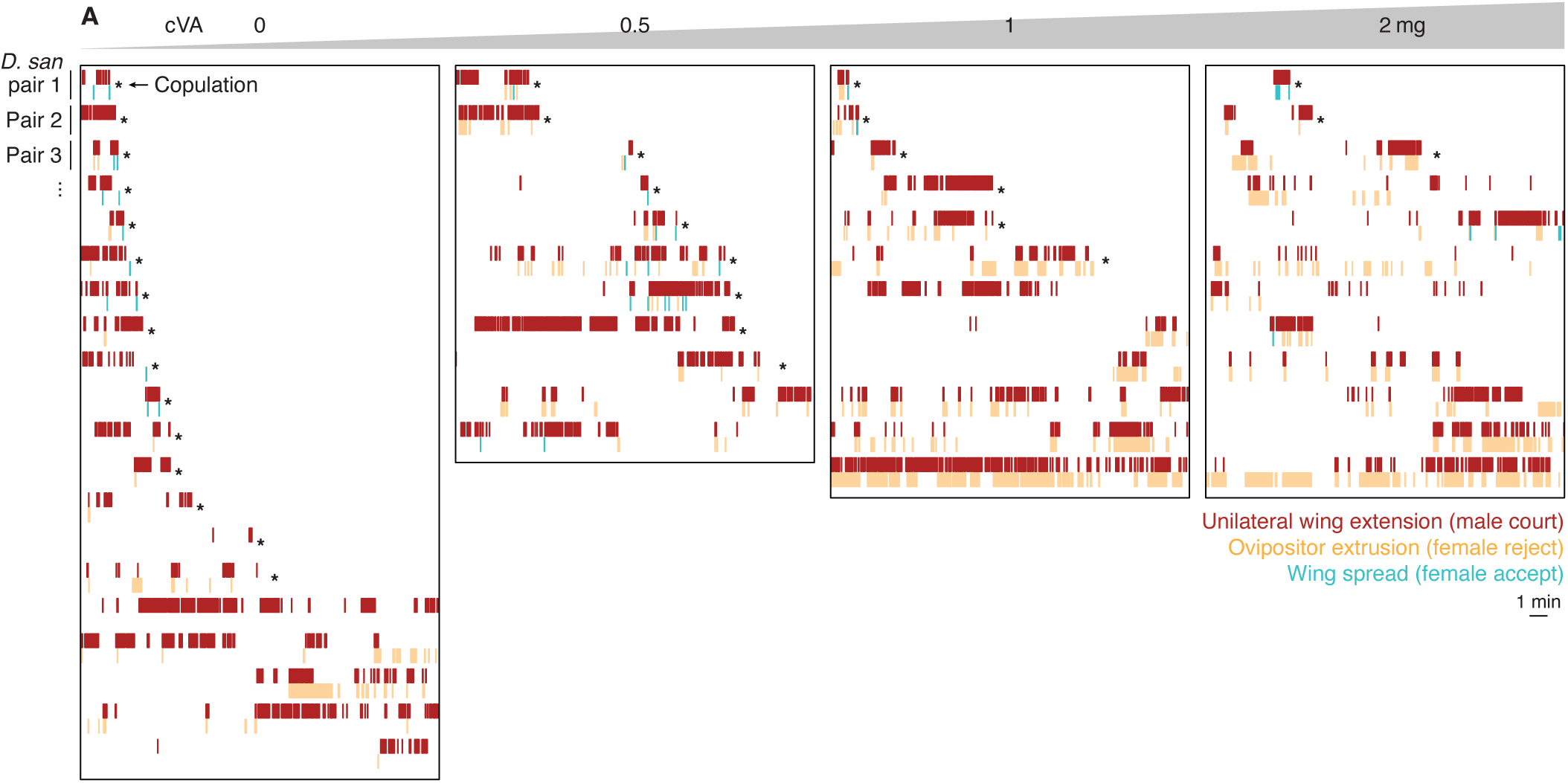
cVA promotes *D. santomea* female sexual rejection during courtship by conspecific males. **(A)** Rasters of *D. santomea* female and male sexual behaviors during dyadic courtship interactions with increasing cVA on a nearby piece of filter paper. For each pair, male courtship (UWE, red) is shown above and female rejection (ovipositor extrusion, orange) or acceptance (wing spreading, cyan) below. Timing of copulations indicated with asterisks. cVA doses indicated at top. Note increased rejection and decreased copulation rates at 1 and 2 mg cVA.

**Figure S18.**
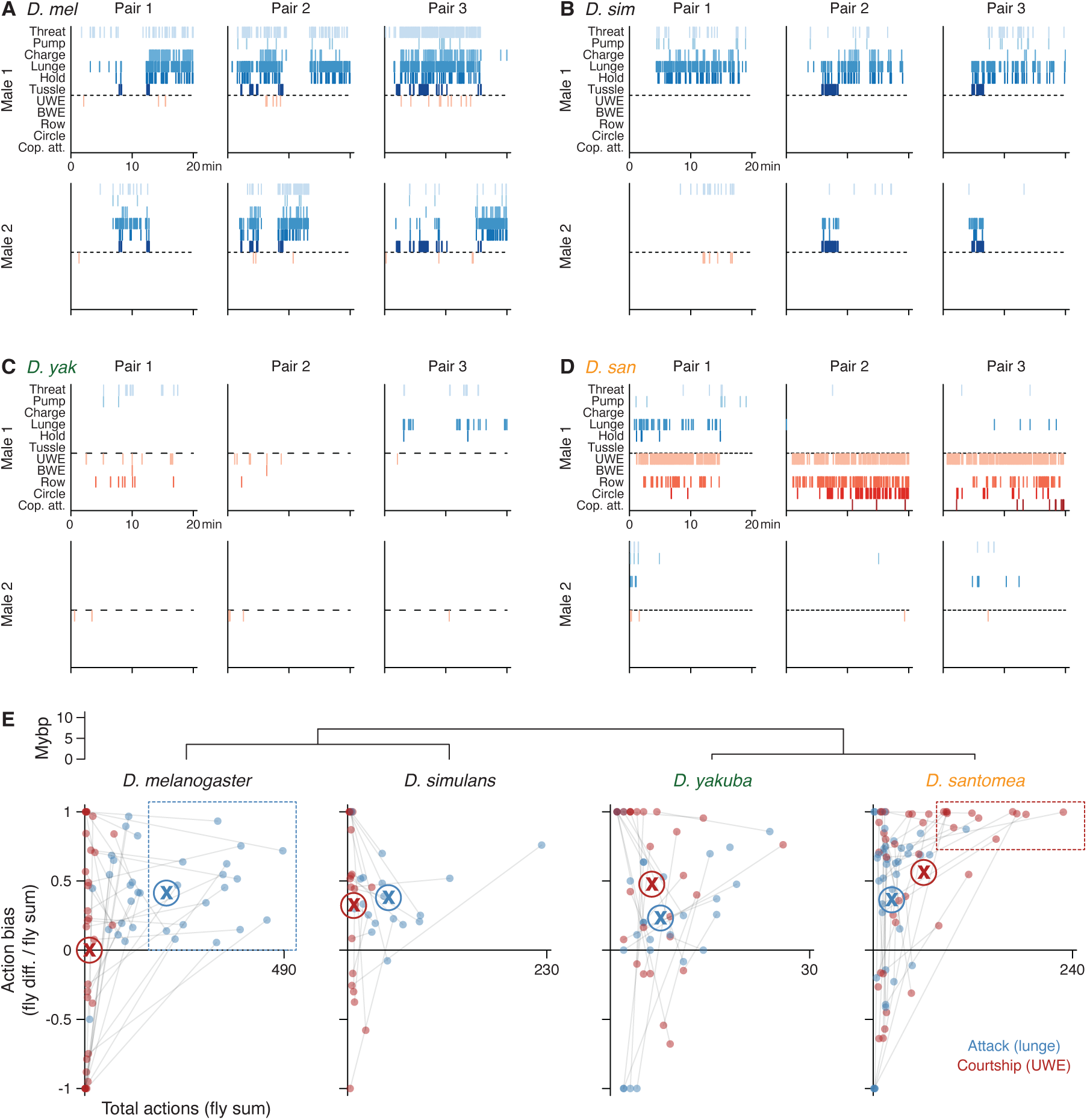
Aggression and courtship biases in *melanogaster* subgroup male-male pairs. **(A-D)** Behavior rasters for three twenty-minute conspecific male-male pairs each of wildtype *D. melanogaster* (A), *D. simulans* (B), *D. yakuba* (C), and *D. santomea* (D) (same pairs as in Fig. 1A). Annotations of six aggressive interactions by manual scoring (blues) are shown above the horizontal dashed lines and five courtship actions (reds) beneath. Actions split by identity of the male to which they are attributed with the one exhibiting the greater total number of actions assigned as “Male 1.” *D. melanogaster* and *D. simulans* males fight with some characteristic bias between flies, *D. yakuba* males are mostly passive but can either fight or court, and *D. santomea* males show frequent and elaborate courtship often with strong bias. **(E)** Spontaneous action biases observed between flies in male-male pairs. Male-directed courtship (red dots) and attack (blue) are plotted as a function of their bout abundance (x-axis) and bias indices (y-axis). Measurements from the same pair connected by thin lines. Cases where the same fly is dominant for both actions show connections confined to the sector above the x-axis. Circled “X” indicates mean value for the corresponding action by color. Boxed regions indicate pairs with high attack expression and wide range of biases in *D. melanogaster* or high courtship expression and consistently strong bias in *D. santomea,* for visual aid.

**Figure S19.**
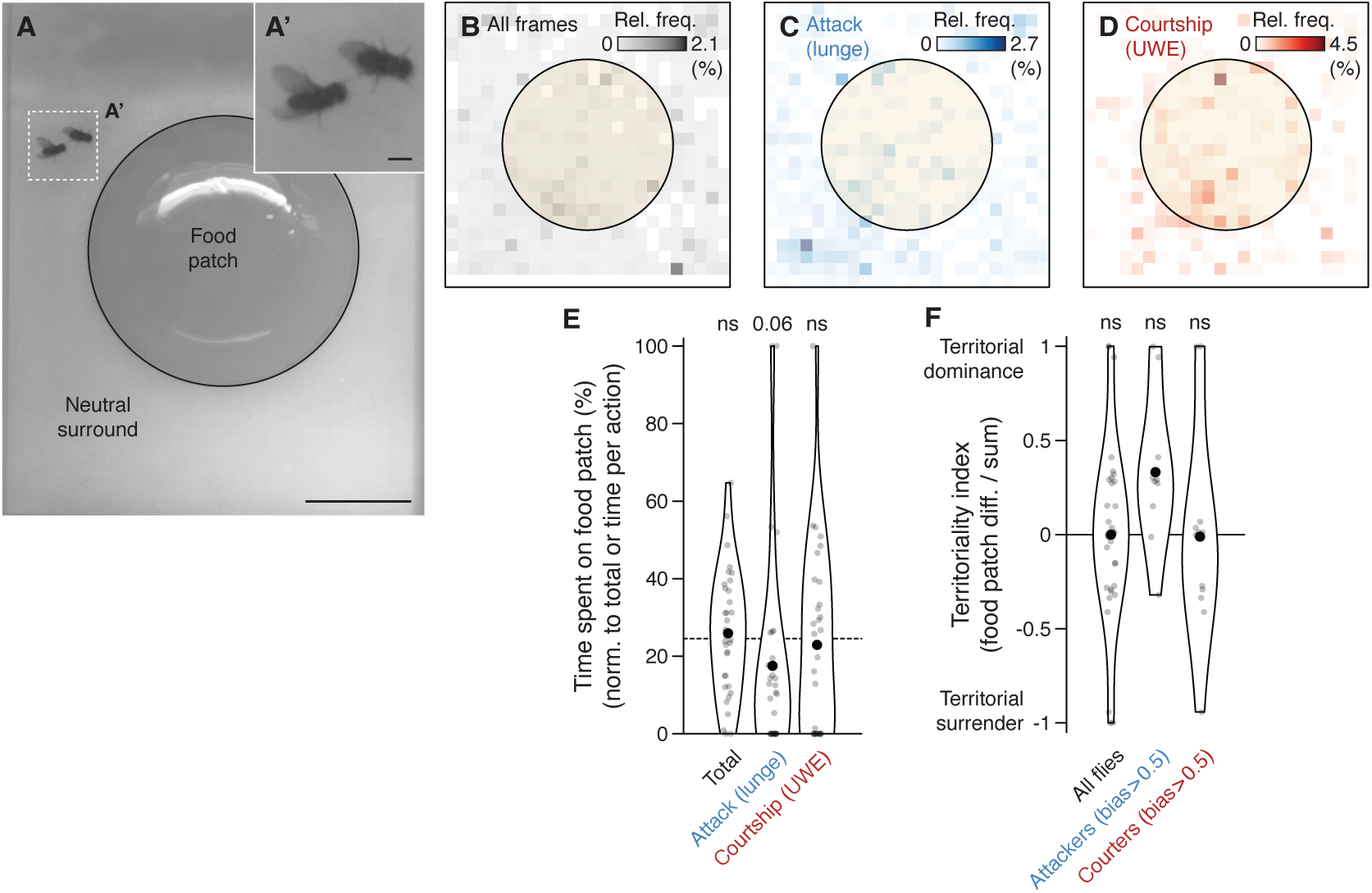
Weak supporting evidence for territoriality in paired *D. santomea* males. **(A)** Video still of a pair of single-housed, wildtype *D. santomea* males in a large behavior chamber (40 mm wide x 50 mm long x 60 mm tall) containing an apple juice-based “food patch” (25 mm diameter) and neutral surround used to test territoriality. (A’) Close-up of intermale courtship from boxed region. Scale bars, 10 mm (A), 1 mm (A’). **(B-D)** Heatmaps representing cumulative spatial patterns of all fly positions (B), attack (lunge) positions (C), and courtship (UWE) positions (D) for 17 *D. santomea* male-male pairs recorded during ten-minute interactions. Normalization in (B) to all frames, in (C) to attack frames only, and in (D) to courtship frames only. Note little sign of increased density on or around the food patch in any case, unlike previously observed for *D. melanogaster* aggression in similar assays (*51*). **(E)** Quantification of (B-D) showing the fraction of time spent on the food patch in total, during attack, and during courtship for each individual *D. santomea* male. Normalizations as in heatmaps above. None are significantly different from having a null median of 25% (random chance, calculated as surface area of the food patch relative to the full arena) by one-sample Mann-Whitney *U* tests. Attack frames show near-significant deviation from chance, but in the opposite direction from that expected for territoriality (enrichment off the food patch). **(F)** “Territoriality index” for *D. santomea* males calculated as the difference in time spent on the food patch between flies normalized by the sum. First distribution represents all pairs, second filtered for pairs where one of the flies showed attack dominance, and third filtered for pairs where one of the flies showed courtship dominance. None are significantly different from having a null median of zero (no territorial advantage) by one-sample Mann-Whitney *U* tests.

**Figure S20.**
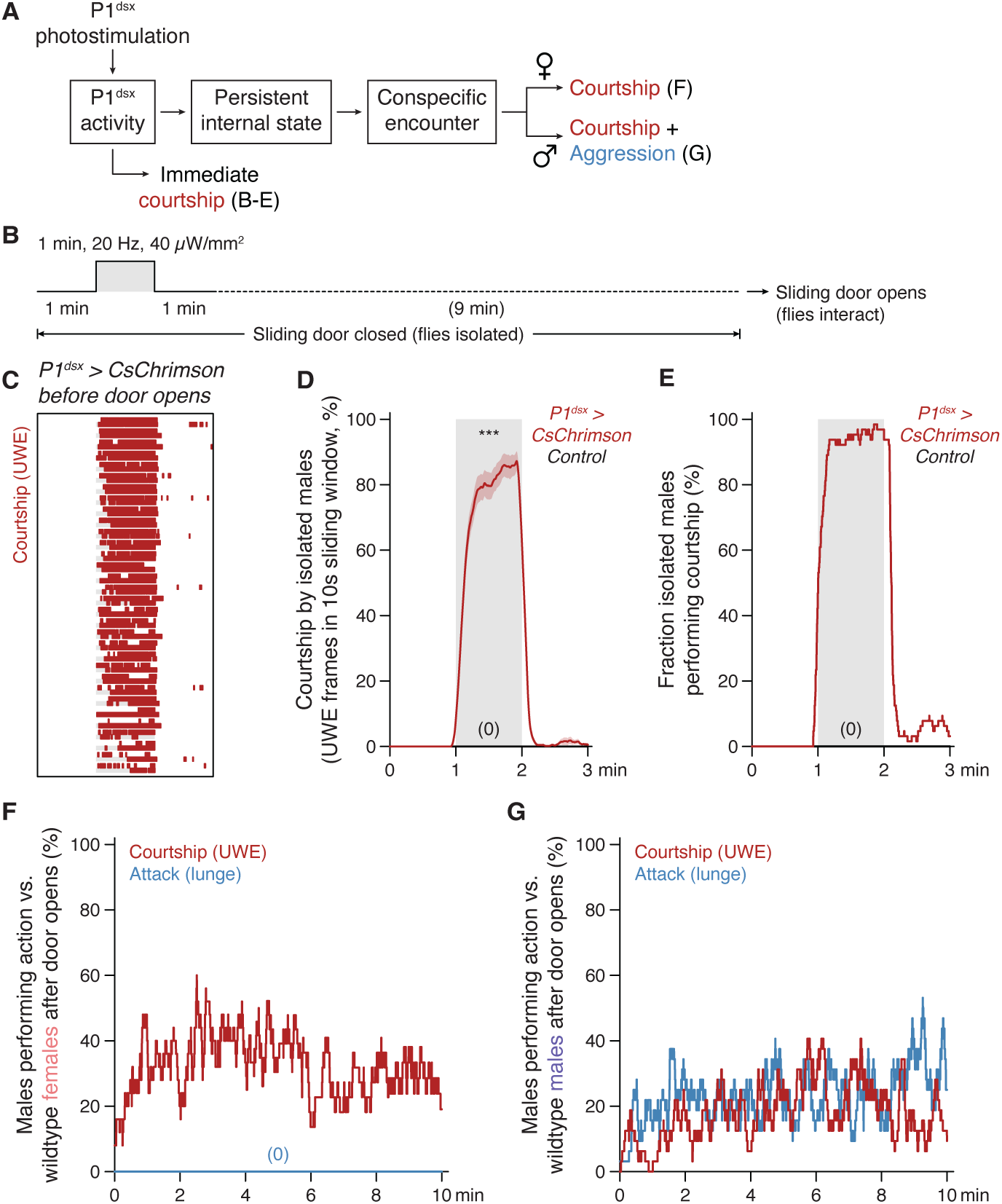
Additional behavior characterizations for the sliding door assay. **(A)** Summary of immediate and delayed behavioral outcomes in *D. santomea* males following brief P1^dsx^ PS in the sliding door assay, with evidence below. P1^dsx^ PS evokes immediate, time-locked courtship during the early isolation phase and in parallel generates a persistent internal state of social arousal. Subsequent encounters with conspecific females or males during the interaction phase elicit pure courtship or mixed courtship and aggression, respectively. Panels containing relevant data for each behavioral outcome are indicated. Figure modified from (*100*). **(B)** Optogenetic photoactivation scheme for isolated *D. santomea* males carrying *P1^dsx^*split-Gal4 (*71G01^DBD^;dsx^AD^*) and Gal4-dependent CsChrimson during the first phase of the delayed encounter (“sliding door”) assay. After a one-minute baseline, two group-housed *P1^dsx^* tester males on either side of a removable divider are exposed to a one-minute photostimulation (PS) block. PS is followed by a ten-minute delay before doors are opened and flies allowed to interact. Delay phase shown as split into one-and nine-minute subphases to reflect cessation of detailed courtship annotations one minute after PS offset. **(C)** Manually scored courtship rasters (UWE, red) for isolated *P1^dsx^* tester males during the first three minutes of the trial. **(D,E)** Expressivity (C) and penetrance (D) of courtship elicited by PS. (C) Traces show the mean fraction of frames containing UWE in a ten-second sliding window with s.e.m. envelopes for *P1^dsx^* (red) and a genetic control (black, no courtship observed). (D) Fraction of flies showing courtship. Note extremely high penetrance and expressivity of induced courtship time-locked to PS in *P1^dsx^*testers. **(F,G)** Fraction of *P1^dsx^* tester males showing courtship (red) and attack (blue) toward wildtype conspecific females (E) or males (F) during the ten-minute interaction phase after doors open. Exclusive courtship is observed toward females and a mix of courtship and attack toward males.

**Figure S21.**
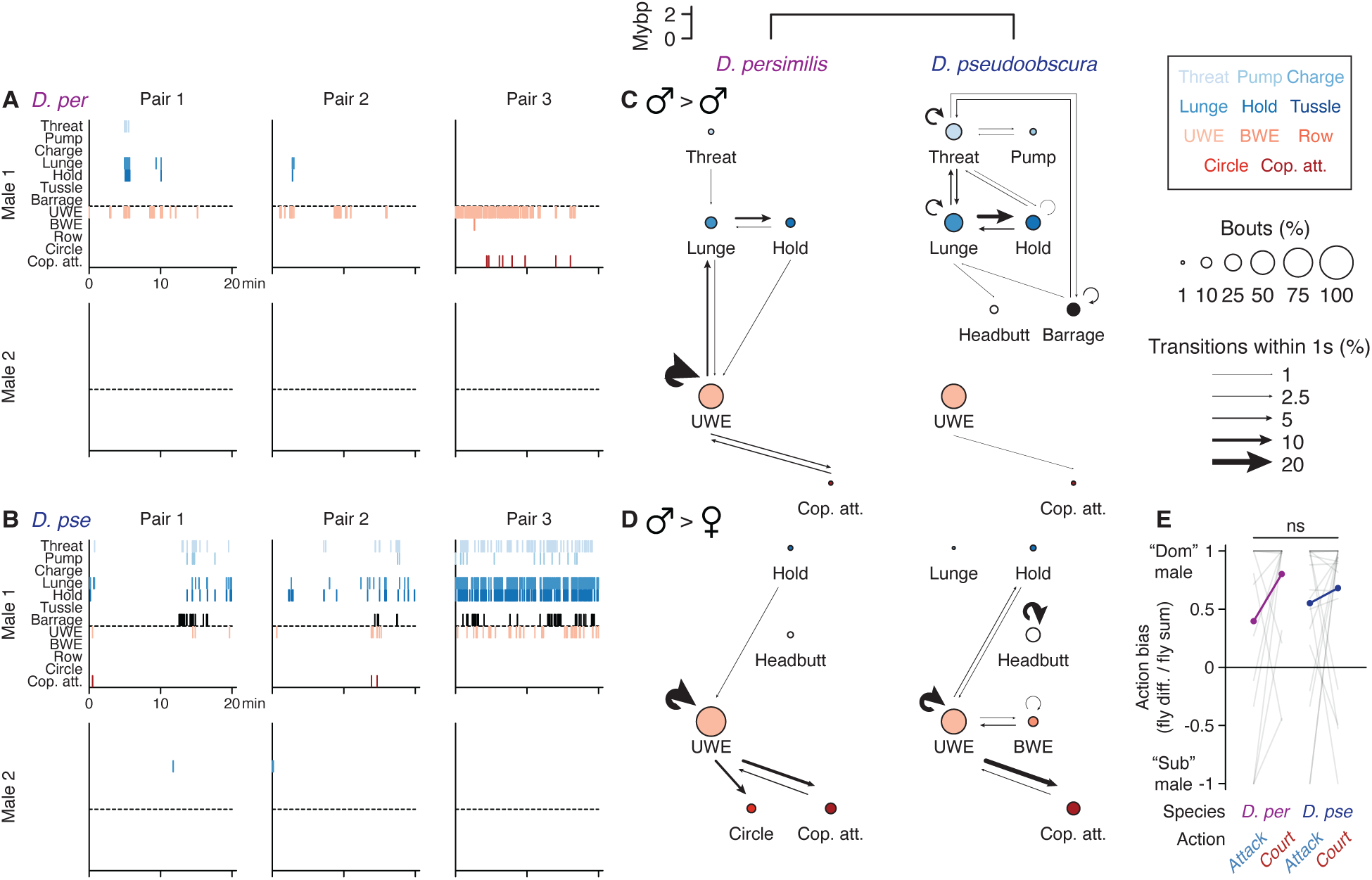
Aggression and courtship by *D. persimilis* and *D. pseudoobscura* males. **(A,B)** Behavior rasters for three twenty-minute conspecific male-male pairs each of wildtype *D. persimilis* (A) and *D. pseudoobscura* (B). Annotations of seven aggressive interactions by manual scoring (blues and black) are shown above the horizontal dashed lines and five courtship actions (reds) beneath. Actions split by identity of the male to which they are attributed with the one exhibiting the greater total number of actions assigned as “Male 1.” *D. persimilis* males court frequently including making copulation attempts (e.g., Pair 3). *D. pseudoobscura* also show some courtship but attack more intensely and often. **(C,D)** Ethograms representing male-(C) and female-directed social behaviors (D) by *D. persimilis* and *D. pseudoobscura* males. Male-female ethograms represent expert annotation for three recordings of male-female pairs per species, with male flies similarly prepared by single-housing and interactions taking place under identical conditions. Male-directed ethograms reproduced from fig. S1A for visual comparison to female-directed counterparts. Note similarity between interactions with males and females in *D. per*. **(E)** Spontaneous bias observed between flies in male-male *D. persimilis* and *D. pseudoobscura* pairs for attack and courtship. Indices derived from the same fly pair are connected as lines and distribution means shown as paired purple (*D. per*) or blue dots (*D. pse*). Both species show strong behavioral biases.

## Materials and Methods

### Fly stocks

All species, stocks, and crosses were reared at 24-25C and 50% humidity on a 12hr:12hr light cycle with standard Caltech fly media. Details about the flies used in each experiment (species, strain, sex, genetic background, and experimental and control genotypes) are listed in table S2 and table S3. Table S2 provides identifiers for the 18 wildtype strains and lists their usage by figure. Table S3 provides similar information for *D. santomea* transgenic lines. All wildtype strains were obtained from the National Drosophila Species Stock Center (Cornell College of Agriculture and Life Science, Ithaca, NY) except *D. melanogaster* Canton-S (Anderson lab, Caltech) and three additional *D. santomea* strains (David Stern lab, Janelia Research Campus).

### Generation of transgenic and knock-in alleles in *D. santomea*

Transgenic *D. santomea* driver and hemidriver lines using enhancer fragments to regulate Gal4 expression were made using phiC31 recombination (*111*) into existing attP landing sites (*112*). White-mutant *D. santomea* attP sites 2251 (2R) and 2253 (3R) (*112*) were targeted using Gal4 (pBPGUw), Gal4DBD (pBPZpGAL4DBDUw), or p65AD (pBPp65ADZpUw) vectors carrying an attB recognition site and mini-white selection marker (*159*, *160*). When available, ready-made constructs with desired enhancer sequences were obtained from Janelia. Otherwise enhancer sequences were obtained from (*159*), synthesized by Twist Bioscience (South San Francisco, CA) with flanking attL1/L2 recognition sites, and cloned upstream of transcriptional control cassettes in the backbone vectors using Gateway LR reactions (Thermo Fisher Scientific, Waltham, MA). Assembled constructs were verified by sequencing and restriction digest, maxi-prepped using endotoxin-free kits, and cleaned using ethanol precipitation before sending for injection. Rainbow Transgenic Flies, Inc. (Camarillo, CA) generated injection mixes containing the construct and phiC31 integrase mRNA (and sometimes also integrase plasmid to increase efficiency) and performed injections into *D. santomea* eggs in batches of 200-250 per desired transformant. Viable G0 (injected) flies were collected and crossed to a white-mutant strain (*2041/w-[3.2]*) (*112*) before screening for yellow-eyed (transgene heterozygous) flies in the G1 progeny to confirm germline transmission. Heterozygotes were crossed together to generate transgene homozygotes as stable lines used for experiments.

Knock-in driver lines were generated using CRISPR/Cas9-mediated homology-directed repair (HDR) (*109*). Donor construct assembly and sgRNA selection for *D. santomea dsx-p65AD* were performed previously (*110*) but transformants positive for the selection marker (*MHC-DsRed*) had not been obtained. The same reagents were re-injected into *D. santomea 146/STO-CAGO 1482* (David Stern, Janelia) in a mixture with nls-Cas9-nls protein (PNA Bio, Thousand Oaks, CA) and one transformant with germline transmission of abdominal mCherry was successfully recovered. Crosses among G1 progeny yielded sporadic G2 flies with the expected *dsx* genital phenotype, verifying the genomic insertion.

*D. santomea Or67d^Gal4^* was generated using the Atalanta strategy (*158*). A backbone vector with two U6 promoters (for expression of two sgRNAs) and an “empty payload” (pJAT32) was selected as the Gateway acceptor. Two donor sequences were synthesized by Twist Bioscience and cloned into the acceptor in a simultaneous double-Gateway reaction. The first (flanked by attL3/L4) is a *T2A-Gal4-stop(hsp70)* cassette (*Gal4-stop(hsp70)* from pBPGUw) designed to be inserted downstream from but as close as possible to the *D. santomea Or67d (3L:LOC120447699)* translational start codon. Placement of the adjacent homology arm ensures that insertion occurs in-frame with the residual 15 coding base pairs so that Gal4 is properly translated. The second (flanked by attL5/L6) is an *ie1-mCherry-stop(p10)* selection marker (*32*) with inverted orientation relative to Gal4. Both donors also contain a 1 kb homology arm and U6-tRNA-sgRNA-tRNA array which, following LR recombination, is positioned downstream of U6. Candidate sgRNAs were designed in Geneious Prime (Auckland, NZ) and a pair was selected which excises 99% of *Or67d* coding sequence, resulting in a null allele (upstream *AATGGCAAAAGTTGAGCCCG*, downstream *ACTGCTGGTTTCCAAAGGAG*). Successful Gateway products were verified by restriction digest and tiled sequencing. Injection into *D. santomea 146/STO-CAGO 1482* was performed by Rainbow Transgenic Flies using nls-Cas9-nls protein as the enzyme source and G1 progeny were screened for germline-transmitting abdominal mCherry. Multiple independent insertion lines were recovered and homozygosed, one of which was used throughout after anatomical verification of uniglomerular labeling in the antennal lobes of the size and position expected for Or67d OSNs (*145*). Note that in this strategy Gal4 expression is controlled exclusively by the *Or67d* native genomic regulatory context rather than an exogenous promoter.

### Spontaneous behavior experiments with video recording

Experimental crosses were seeded with 10-12 virgin females and 5-6 males and flipped every 3-4 days. Experimental flies were collected mostly as virgins on the day of eclosion and reared either in isolation (single-housed, 1 individual per vial) or in same-sex groups (group-housed, 10-20 individuals per vial) for 5-6 days until testing. Group-housed (but not single-housed) flies were flipped into fresh vials 1-2 days before testing. For experiments requiring unambiguous determination of individual identity (e.g., in mixed-genotype or mixed-species male-male pairs), a small distal portion of one fly’s wing was clipped using a razor blade under light anesthesia. Whenever possible, the “target” fly (individual whose behavior was not being measured) was selected for wing clipping. Otherwise clips were balanced between the two interacting species or genotypes. Females were collected and group-housed as were males (mostly as virgins on the day of eclosion, 10-20 individuals per vial) and age-matched to the males they were tested with as courtship targets. As this was usually a period of 5-6 days, it is likely that even the few females that were not virgins upon collection had by the time of the experiment shed residual male pheromones potentially transferred during prior copulation (e.g., cVA). Group-housed females were also flipped 1-2 days before testing. Housing for each experiment arranged by figure can be found in table S2 and table S3 (“single” or “group” under the “Housing” heading).

Behavior experiments took place in the morning between 9:00AM (ZT0) and 1:00PM (ZT4) in a temperature and humidity-controlled room (24C, 50% humidity) with ambient white lighting. Single, pairs, or trios of flies interacted in 8-well acrylic circular chambers (∼2.0 cm^2^ surface area, ∼2.4 cm^3^ volume) designed and described previously (*57*, *161*). Walls of the chambers were coated with Fluon (Tar Heel Ants, Raleigh, NC) and lids with a siliconizing fluid (Thermo Fisher) to discourage flies from climbing. Chamber floors were covered with a thin layer of freshly prepared “food” (2.5% (w/v) sucrose, 2.25% (w/v) agarose in Tropicana apple juice) and illuminated with an 855 nm backlight (Smart Vision Lights, Muskegon, MI). Experiments testing alternative foods replaced the apple with other juices acquired commercially (*Morinda*, fig) or from botanical gardens (*Marula*, *Pandanus*), or with water as a “no food” condition (sucrose was also omitted in this case). Flies were introduced one at a time into chambers by gentle aspiration through a small hole in the lid. Interactions were video recorded from above using a Point Grey Flea3 camera running at 30 Hz with a 780 nm long pass IR filter (Midwest Optical Systems, Palatine, IL). Experiments performed in 8-well chambers are listed as “small” under the “Chamber” heading in table S2 and table S3.

For copulation tests of Or67d OSN-silenced *D. santomea* females and territoriality assays of wildtype *D. santomea* males, “large” chambers consisting of a central food patch (apple juice agarose) and a neutral surround (bare acrylic) were used. Chambers measure 20 cm^2^ surface area and 120 cm^3^ volume, and the circular food patch ∼4.9 cm^2^ (25% surface area). Flies (12 females and 12 males for copulation tests, 2 males for territoriality tests) were gently aspirated into chambers and video recorded using identical lighting and camera settings as for small chambers described above.

### Optogenetic photoactivation experiments

Flies were collected on the day of eclosion into vials containing food supplemented with 400 µM all *trans*-Retinal (Millipore Sigma, Burlington, MA) and reared in the dark for 5-6 days, with one flip into fresh supplemented vials 1-2 days before testing. Photoactivated flies were usually group-housed (10-20 males per vial), except in male Or67d OSN courtship suppression experiments which were single-housed to encourage spontaneous courtship prior to stimulation. Group-housed males paired together during behavior assays came from different vials.

Experiments took place in the small chambers using an 8-LED array setup reported previously (*57*, *161*). LEDs delivered flashing 685 nm red light in photostimulation (PS) blocks with 10 ms pulse width, frequencies ranging from 1-40 Hz, and intensities 10-260 µW/mm^2^, calibrated with a photodiode power sensor (S130VC, Thorlabs, Newton, NJ) placed at the location of the behavior chambers. Exact PS conditions (timing, frequency, intensity) for each experiment are shown in Figures and described in Figure Legends.

“Sliding door” experiments testing persistent behavioral responses after PS were performed in 8-well chambers fitted with vertical slits to allow metal barriers to divide each chamber in half. Barriers remained in place during the isolation/PS phase of the experiment, which upon completion of the delay were manually removed during a ∼30 second period while video recordings were paused. Once the barriers were removed recordings resumed for the interaction phase.

### Behavior annotations and analysis

Annotations of aggressive (A) and courtship behaviors (C) in male-male pairs from each species for the primary screen were performed manually using action definitions from (*56*) (threat (A), charge (A), lunge (A), hold (A), tussle (A), unilateral wing extension (C), circle (C), copulation attempt (C)), (*106*) (pump (A)), (*20*) (bilateral wing extension (C), row (C)), and (*157*) (headbutt (A)). One additional action observed exclusively in *D. pseudoobscura* males, “barrage,” is defined as follows: *the attacking fly pushes into and quickly chases the opponent while both wings are fully extended and vibrating rapidly as if in flight.* This action appears to have been described once before without being named (*136*). Ethograms were constructed from calculations of action frequencies and transition matrices using a 1-second cutoff.

For most experiments, video recordings were analyzed in an automated fashion by first tracking the absolute and relative positions of flies in the chamber at every frame (Caltech FlyTracker, https://kristinbranson.github.io/FlyTracker/) (*56*) and using the tracking information as input to automated behavior classifiers developed previously for *D. melanogaster* social interactions (*57*), implemented in JAABA (https://jaaba.sourceforge.net/) (*55*). Two classifiers were used extensively after satisfactory performance evaluation for species within the melanogaster subgroup: “lunge” for consummatory, contact-mediated male aggression, and “unilateral wing extension” for male courtship. Recordings analyzed in this way are listed as “automated” under the “Behavioral analysis” heading in table S2 and table S3. Manually annotated recordings, including ones analyzed for the female sexual behaviors ovipositor extrusion (*126*, *127*) and wing spreading (*128*), are instead labeled “manual.” For experiments involving a male-female pair, analysis ended at the point of copulation (if it occurred) and the latency to copulation was used for normalization procedures rather than total interaction time.

### Courtship song recordings and analysis

Flies (*D. santomea* wildtype strain 14021-0271.00) were raised in standard laboratory conditions at 23C and 12:12 light cycle. Sexually naïve individuals were isolated within 3 hours of eclosion and single-housed for 5-7 days prior to behavioral analysis. Synchronized audio and video song recordings were made within 4 hours of lights-on in the morning using the Song Torrent system (*60*). Individual flies arranged in male-male or male-female pairs were loaded onto opposite sides of a divider in recording chambers and introduced to each other simultaneously at the beginning of 30-minute recordings.

Videos of 16 male-male and 16 male-female pairs of *D. santomea* flies were visually scanned and 3 pairs of each type with ample courtship were selected for audio analysis. Song traces were visualized and annotated in Tempo (*63*) until dozens to hundreds of both “pulse” and “clack” events had been recorded for each courting male (instead of to saturation). The exact amplitude and position of each event’s peak was determined by obtaining the maximum signal intensity within a 20 ms time window centered on the annotation point. Events were assigned to being part of a “train” when their separation from another event of the same type was less than 200 ms for pulse and 400 ms for clack.

### Pheromone applications

(Z)-11-octadecenyl acetate (cVA) and (Z)-7-tricosene (7-T) were acquired commercially from Pherobank B.V. (Wijk bij Duurstede, NL) as neat liquid formulations (i.e., without a solvent). For behavioral experiments with cVA on filter paper, 0.5-2 µl neat formulation was applied just prior to introduction of flies into the chamber. For direct treatment of live flies, cVA neat formulation was diluted to a 10% working solution in acetone (179124, Millipore Sigma), from which 0.2 µl was pipetted onto the abdomen of target flies under anesthesia 1-2 hours prior to behavioral testing, as previously (*89*). Flies that did not rouse were discarded. For direct treatment of dead flies, 0.2 µl of 10% cVA (acetone) or 7-T (hexane) working solutions was directly pipetted onto the abdomen of flies freshly killed by freezing at −80C and glued to the chamber floor using melted agarose of the same “food” composition.

### Odor place preference tests

The olfactory “y-maze” was constructed by laser cutting multiple layers of 1/8th inch 3143 infrared (IR)-transmitting acrylic (ACRY31430.125PM, ePlastics, San Diego, CA) to shape. The bottom floor layer was finely wet sanded for texture. Chambers at the end of each arm meant for odor reservoirs and a center port for the vacuum line were blocked off with fine mesh. Top layers consisted of a ceiling of static dissipating acrylic (8774K11, McMaster-Carr, Santa Fe Springs, CA) with a rim of IR acrylic to block ambient light, and a final roof layer with small holes above the chambers to allow air to be drawn into and down the arena by the vacuum line. The layers were supported and sealed with three threaded rods and bolts. Acrylic layers were periodically cleaned with 1% SDS, water, and isopropanol. The arena was illuminated from below using a custom-built IR 810 nm LED lighting board and diffused with semi-opaque white acrylic. Videos were collected with a machine vision camera (BFS-U3-04S2M-C: 0.4 MP, Teledyne FLIR) running at 45 Hz and a Pentax 12mm 1:1.2 TV lens (FL-HC1212B-VG, Ricoh).

Flies were collected on the day of eclosion, separated by sex, and single- or group-housed for one week until testing. For each trial, 5 µl odor (10% cVA in acetone or acetone control) was first loaded at the top of each arm of the y-maze then 2-4 cold anesthetized males or females were aspirated into the central bowl and the lid quickly fastened. Ten-minute recordings initiated once the flies had roused. Cumulative position traces were obtained by first median filtering on a subset of frames to construct a background image. OpenCV (4.6.0) was used to perform thresholding on background-subtracted frames. The resulting “blob detection” indicated animal position per frame, and positions across the entire trial were plotted using Matplotlib (3.4.3). Normalized occupancy in the left vs. right arm was calculated from cumulative position traces using cutoffs including the two arms but excluding the central bowl.

### Gas chromatography – mass spectrometry (GC-MS)

Flies were collected on the day of eclosion, separated by sex, and group-housed in sets of 10 for one week until analysis. Flies were anesthetized and transferred to 2 ml glass screw top vials (Thermo Fisher) containing 100 µl hexane (139386, Millipore Sigma) with 10 ng/µl octadecane (C18) internal standard (O652, Millipore Sigma). Hexane extraction was allowed to proceed for 15 minutes at room temperature, after which extracts were transferred to glass inserts (Ibis Scientific, Clark County, NV) positioned in new screw top vials for either immediate analysis or storage at −80C.

An AOC-2i autoinjector (Shimadzu, Kyoto, Japan) was used to inject 2 µl of each sample into a Shimadzu QP-2020 GC-MS system (Shimadzu, Kyoto, Japan) equipped with a ZB-5MS fused silica capillary column (30 m x 0.25 mmID, df=0.25) (Phenomenex, Torrance, CA) and helium as the carrier gas. The injection port was maintained at a temperature of 310C and was operated in splitless mode, with a column flow rate of 2.15 ml/min. The temperature program consisted of a 40C hold for 1 minute, followed by a 20C/min ramp to 250C, a 5C/min ramp to 320C, and a 320C hold for 7.5 minutes. The transfer line to the mass spectrometer was held at 320C and the ion source temperature at 230C. Electron ionization was performed at 70 eV and MS scans were collected at 2 Hz, spanning 40-650 m/z. Identification of cuticular hydrocarbon (CHC) and ester compounds was determined by retention index, diagnostic ions, and comparison to previous GC-MS results from *Drosophila* (*65*, *66*, *162*, *163*). GC-MS spectra were integrated in python using the pyteomics package (v4.4.0).

### Immunostaining and imaging

Whole brains were dissected in ice-cold phosphate-buffered saline (PBS) and fixed in 4% paraformaldehyde for 20 minutes at room temperature (RT). Samples were washed three times in 0.5% PBST (PBS containing 0.5% (v/v) Triton X-100) for 10 minutes at RT, then permeabilized in 2% PBST for 30 minutes at RT. Samples were blocked in 0.5% PBST with 5% normal goat serum (NGS) for 1 hour at RT or overnight at 4C. Samples were incubated with primary antibodies in blocking solution for 48-72 hours at 4C (rabbit anti-DsRed 1:1000, mouse anti-Bruchpilot 1:50) followed by three 10-minute washes in 0.5% PBST. Samples were incubated in secondary antibodies (anti-rabbit IgG AF568 and anti-mouse IgG AF633, 1:1000) in blocking solution for 36-48 hours at 4C, then again washed three times in 0.5% PBST for 10 minutes at RT before mounting.

Samples were mounted in VectaShield antifade medium (Vector Laboratories, Newark, CA) and imaged on an Olympus Fluoview FV3000 confocal microscope (Tokyo, JP) with inverted 30X/1.05 or 60X/1.30 oil immersion objectives. Image stacks were acquired at 1024 x 1024 px^2^ spatial resolution with 1.0-1.5 µm z-step intervals. Images (referred to as “photomicrographs”) are maximum intensity projections (MIPs) of image stacks generated in Fiji. Fiji was also used to pseudocolor, overlay, adjust brightness and contrast, and add scale bars to images.

### Quantification and statistical analysis

Fly positional tracking, automated social behavior annotations, and song analyses were performed using software implemented in MATLAB (MathWorks, Natick, MA). Odor place preference and GC-MS experiments were analyzed in python using Jupyter Notebooks (https://jupyter.org/). The tabulated outputs of these analyses were saved and exported as text files for final analyses in R (R Project for Statistical Computing) using RStudio (Posit PBC, Boston, MA). Custom scripts in R were used to calculate raw data derivations reported in figures, produce plots visualizing them, and perform statistical tests. Illustrator 2025 (Adobe, San Jose, CA) was used to adjust aesthetic features of plots and assemble figures.

Measurement distributions are often shown as boxplots highlighting the median and overlayed data points. Outliers are identified in the standard way (if > Q3+1.5*QR or if < Q1-1.5*QR) and visually excluded from box whiskers, but are retained in all quantifications (including statistical tests) and still shown as overlayed dots. Violin plots show full distribution ranges as vertically mirrored kernel density plots with mean highlighted. Statistical tests consisted mostly of non-parametric comparisons between pairs of or multiple distributions and were implemented in R. Mann-Whitney *U* tests (R base *stats* package: *wilcox.test* function) were used to compare pairs of distributions (two-sample) or an empirical distribution to a null median (one-sample), and could be paired or unpaired depending on whether the comparison involved the same individual or interacting pair. Two-sample Kolmogorov–Smirnov tests (*stats: ks.test*) were used to compare pairs of time-evolving distributions or cumulative distributions. Kruskal-Wallis tests (*stats: kruskal.test*) and post-hoc Dunn’s tests (*FSA: dunnTest*) were used to compare multiple distributions and assign lettered groupings. Binomial tests (*stats: binom.test*) were used to compare pairs of frequencies for binary outcomes (two-sample) or an empirical frequency to a null probability (one-sample). Linear fits and Pearson correlations were calculated using the *lm* and *cor* functions (*stats*), respectively. Benjamini-Hochberg multiple test correction was used to adjust p-values whenever appropriate (*stats: p.adjust: method=‘BH’*). Standardized effect sizes were calculated using Cohen’s *d* (*effsize: cohen.d*). Tests shown in each figure can be referenced in table S4 for details including sample sizes, sample units, test used, whether correction was applied, adjusted p-value, and effect size. Significance levels are reported in Figures as *<0.05, **<0.01, and ***<0.001.

## Notes

### Competing Interest Statement

The authors have declared no competing interest.

### Summary of Updates

Minor edits were made to text and figures

